# The effect of single mutations in Zika virus envelope on escape from broadly neutralizing antibodies

**DOI:** 10.1101/2023.09.13.557606

**Authors:** Caroline Kikawa, Catiana H. Cartwright-Acar, Jackson B. Stuart, Maya Contreras, Lisa M. Levoir, Matthew J. Evans, Jesse D. Bloom, Leslie Goo

## Abstract

Zika virus and dengue virus are co-circulating flaviviruses with a widespread endemic range. Eliciting broad and potent neutralizing antibodies is an attractive goal for developing a vaccine to simultaneously protect against these viruses. However, the capacity of viral mutations to confer escape from broadly neutralizing antibodies remains undescribed, due in part to limited throughput and scope of traditional approaches. Here, we use deep mutational scanning to map how all possible single amino acid mutations in Zika virus envelope protein affect neutralization by antibodies of varying breadth and potency. While all antibodies selected viral escape mutations, the mutations selected by broadly neutralizing antibodies conferred less escape relative to those selected by narrow, virus-specific antibodies. Surprisingly, even for broadly neutralizing antibodies with similar binding footprints, different single mutations led to escape, indicating distinct functional requirements for neutralization not captured by existing structures. Additionally, the antigenic effects of mutations selected by broadly neutralizing antibodies were conserved across divergent, albeit related, flaviviruses. Our approach identifies residues critical for antibody neutralization, thus comprehensively defining the as-yet-unknown functional epitopes of antibodies with clinical potential.

**Importance:** The wide endemic range of mosquito-vectored flaviviruses – such as Zika virus and dengue virus serotypes 1-4 – places hundreds of millions of people at risk of infection every year. Despite this, there are no widely available vaccines, and treatment of severe cases is limited to supportive care. An avenue towards development of more widely applicable vaccines and targeted therapies is the characterization of monoclonal antibodies that broadly neutralize all these viruses. Here, we measure how single amino acid mutations in viral envelope protein affect neutralizing antibodies with both broad and narrow specificities. We find that broadly neutralizing antibodies with potential as vaccine prototypes or biological therapeutics are quantifiably more difficult to escape than narrow, virus-specific neutralizing antibodies.

## Introduction

Viruses can evade immune pressures by antigenic variation and adaptation. For co-circulating, antigenically similar viruses, this can mean population immunity to one virus might determine the fitness of another virus strain or serotype. Zika and dengue are two such antigenically related, co-circulating flaviviruses, with dengue virus clustering into four unique serotypes and Zika virus occupying a single serotype^1^. An initial natural infection with any one of these viruses typically elicits a narrow, virus-specific neutralizing antibody response^2^, and, to some extent, this prior exposure dictates risk of future infection with particular strains and serotypes^3,4^. Another consequence of this effect at the population level is that dengue virus outbreaks are measurably strain- and serotype-dependent, and recur in waves consistent with patterns of immune evasion^5,6^. These dynamics are also complicated by evidence of considerable genetic and antigenic diversity within Zika virus and dengue virus serotypes^2,3,7–9^.

Furthermore, primary exposure to Zika or dengue virus has also been shown to increase risk of severe disease upon a second infection with a heterotypic dengue virus^10–12^. The widely accepted explanation for this increased risk is antibody-dependent enhancement, wherein viruses complexed with cross-binding but non-neutralizing antibodies are internalized into cells expressing Fc receptors^13–15^. Thus, there is immense interest in both developing vaccines that elicit broad and potent neutralizing antibody responses, as well as isolating and leveraging broadly neutralizing antibodies as biological therapeutics. Towards this end, investigations have examined human antibody repertoires and isolated a select few antibodies that can broadly and potently neutralize Zika virus and all serotypes of dengue virus^16–20^. These antibodies all target envelope (E) protein, which is the major antigenic target in both Zika virus and dengue virus^21,22^. Due to their therapeutic potential, much effort has gone towards further characterization of the structure and binding requirements of these antibodies^20,23–28^.

While proposing these broadly neutralizing anti-E antibodies as therapeutics or as a template for vaccine design^29,30^ is potentially exciting, important biophysical and evolutionary questions remain. In these instances, the immune system has managed to generate antibodies that cover considerable E protein antigenic diversity—but what is the effect of further viral diversification on antibody neutralization? It is well-established that only a handful of the residues identified as an antibody’s binding footprint (ie., the structural epitope) are functionally required for neutralization (ie, the functional epitope)^31^. It would therefore be valuable to know exactly which mutations would affect neutralization by broadly neutralizing anti-E antibodies.

Here, we use deep mutational scanning to identify functional epitopes by measuring the effect of all possible single amino acid mutations in Zika virus E protein on neutralization by both broadly neutralizing and narrow virus-specific antibodies. We find that while all antibodies are affected by single amino acid mutations, the magnitudes of these effects vary widely across antibodies. Specifically, single mutations only modestly increase neutralization resistance against broad antibodies, while single mutations completely ablate neutralization by antibodies with narrow specificities. We extrapolate these results to other flavivirus genetic contexts by engineering neutralization escape mutations in other Zika virus and dengue virus E proteins. Interestingly, we show that some antigenic effects are conserved across these overall divergent viral surface proteins.

## Results

### Measuring the effects of all Zika virus E protein mutations on broad and narrow antibody neutralization

We first assembled a panel of neutralizing antibodies of varying breadth and potency against Zika and dengue viruses. The neutralization profiles of these antibodies and their structures in complex with E protein homodimer are shown in **Figure 1**. The antibodies chosen have either quaternary or tertiary epitopes, defined by their ability (or lack thereof) to bind single E protein homodimers, as shown in **Figure 1A**. The most broad and potent of these antibodies belong to the E dimer epitope 1 (EDE1) sub-class, and are characterized by their quaternary epitope spanning both E monomers within the dimer subunit. This intra-dimer epitope includes parts of the fusion loop, a highly conserved region necessary for membrane fusion, and the b loop of domain II^24,26,28^. Both EDE1 subclass antibodies tested, EDE1-C10 and EDE1-C8, neutralize Zika virus and all serotypes of dengue virus^18,27^ (**Figure 1B**). SIgN-3C is an equally broad neutralizing antibody^32^, but unlike EDE1, targets an epitope targeting multiple E dimers^23^. In contrast to EDE1 and SIgN-3C antibodies, MZ4 neutralizes Zika and only two serotypes of dengue virus, and binds to the domain I/II linker within the E monomer^20^. As a control, we profiled the antibody ZV-67, which has narrow, Zika virus-specific activity and targets an tertiary epitope in domain III^33^.

**Figure 1.**
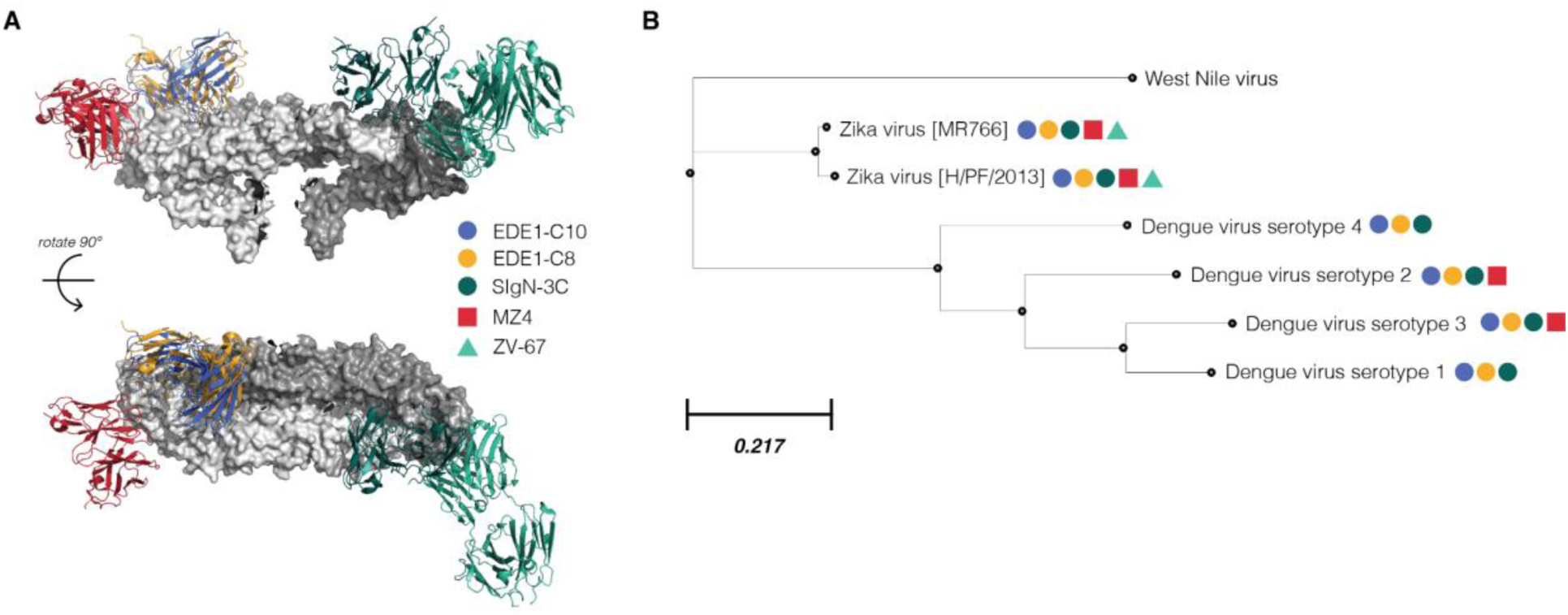
Structural epitopes and breadth of flavivirus-neutralizing antibodies. **(A)** Structures of Zika virus-targeted and broadly neutralizing monoclonal antibodies complexed onto E protein dimer (PDB 5IRE^44^), shown from a side and top-down view. The homodimer is colored light gray and dark gray to distinguish component monomers, and antibodies are colored per the corresponding key. SIgN-3C (PDB 7BUA^23^) targets a complex quaternary epitope, spanning multiple adjacent E homodimers. EDE1-C8 (PDB 5LBS^28^) and EDE1-C10 (PDB 5H37^24^) target a quaternary epitope spanning both monomers within the homodimer. MZ4 (PDB 6NIU^20^) targets a DI-DIII linker region within E monomer. ZV-67 (PDB 5KVG^33^) targets a DIII epitope within the E monomer. **(B)** An inferred phylogenetic tree of E protein sequences from a panel of flaviviruses. Branch lengths are determined by amino acid substitutions per site. Colored dots refer to the same key as in panel A, and indicate that a particular virus is neutralized by the referenced antibody. For instance, ZV-67 only neutralizes Zika virus, whereas EDE1-C10 neutralizes all Zika and dengue viruses.

To perform deep mutational scanning, we regenerated previously described mutant E virus libraries from Zika virus ‘African’ lineage strain MR766^34^. Despite the lab-adapted history of this strain, we reasoned that because Zika viruses share a single serotype^1^, we would be able to identify mutations that affect contemporary strains of Zika virus. We then infected these mutant libraries into cells with each antibody in our panel to select for escape variants, and read out the results by deep sequencing as described previously^34^. Each deep mutational scanning experiment was carried out in biological triplicate using three independently generated mutant libraries at antibody concentrations that neutralized >99% of the mutant virus libraries (**Figure 2A**; **Supplemental Table 1**). Similar to previous work, we reasoned that this would select for escape mutations with the largest individual effects^35^. For the antibody SIgN-3C, we also performed selections at an additional concentration to improve accuracy of identified escape mutations. To quantify mutation-level antibody escape, we calculated the ratio of a given amino acid mutation in E protein in the antibody-selected libraries relative to the unselected libraries. The correlations for replicate selections are shown in **Supplemental Figure 1**.

**Figure 2.**
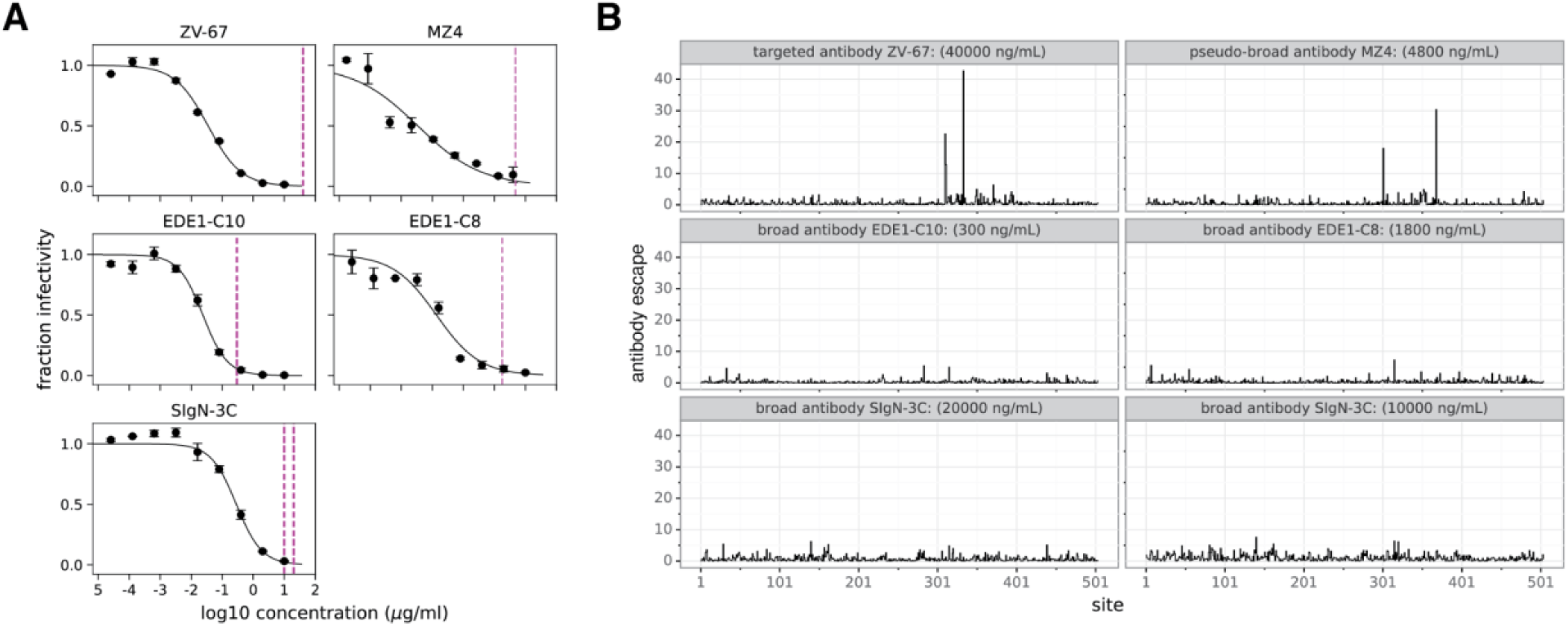
Antigenic effect size across antibody selections at site-level. (**A)** Wild-type MR766 Zika virus neutralized by antibodies indicated above each graph. Dotted lines indicate the antibody concentrations used to neutralize virus libraries for deep mutational scanning. Concentrations were chosen that neutralized >99% of virus libraries. **(B)** The site-wise summed escape from each antibody across the entirety of E protein. For narrow and pseudo-broad antibodies, ZV-67 and MZ4, there are large peaks corresponding to sites of strong antibody escape. For broad antibodies, EDE1-C10, EDE1-C8 and SIgN-3C, there are no single large peaks; instead, there are multiple smaller peaks. The mutation-level antibody escape values are reported in **Supplementary Figures 2-6**. For details on calculation of antibody escape, see **Methods**.

We next plotted the site-level effects of all mutations by summing all individual amino acid antibody escape measurements at a given site in **Figure 2B**. These plots show that there are sites of strong escape from the narrow and pseudo-broad antibodies ZV-67 and MZ4, but only sites of weaker escape from the broad antibodies EDE1-C10, EDE1-C8 and SIgN-3C. For ZV-67 and MZ4, sites with the largest antibody escape occurred within a single E protein region (domain III). In contrast, EDE1-C10, EDE1-C8 and SIgN-3C had smaller peaks of escape scattered across the entire E protein.

The effects of individual amino acid mutations at each site are shown in **Supplemental Figures 2-6**. For ZV-67 and MZ4, many different amino-acid mutations tended to cause antibody escape at the key sites. However, for the broad antibodies typically only one or a handful of mutations at a given site led to viral escape, and the magnitude of escape tended to be lower. A likely explanation is that our approach only identifies escape mutations that are functionally tolerated in live replicative virus, and the broad antibodies tend to target sites in E that are relatively intolerant of mutations.

### Comparing functional epitopes identified by deep mutational scanning to structural epitopes

We next compared the sites of antibody escape from the deep mutational scanning to the structural epitopes of the antibodies as determined by prior structural studies (**Figure 3**). The Zika-specific neutralizing antibody, ZV-67, exhibited antibody escape at only three sites— residues A310, A311 and A333—that comprise the core of the structural epitope^33^ (**Figure 3A**). Particularly at site A333, there were many individual mutations with large antibody escape values, representing amino acids with all types of side chains (hydrophobic, hydrophilic, aromatic, acidic, basic and polar uncharged). In short, a small number of sites comprise the functional epitope of ZV-67; at each of these sites, a diverse array of mutations lead to viral escape. In a similar vein, antibody escape from the pseudo-broad antibody MZ4 is dominated by two sites, K301 and S368 (**Figure 3B**) within the previously described structural epitope^20^. At site K301, deep mutational scanning indicated a wide variety of mutations lead to escape, whereas at site S368 antibody escape was biased towards non-polar hydrophobic mutations. In sum, narrow and pseudo-broad antibodies ZV-67 and MZ4 had a wide variety of escape mutations at a few sites within the structural epitope.

**Figure 3.**
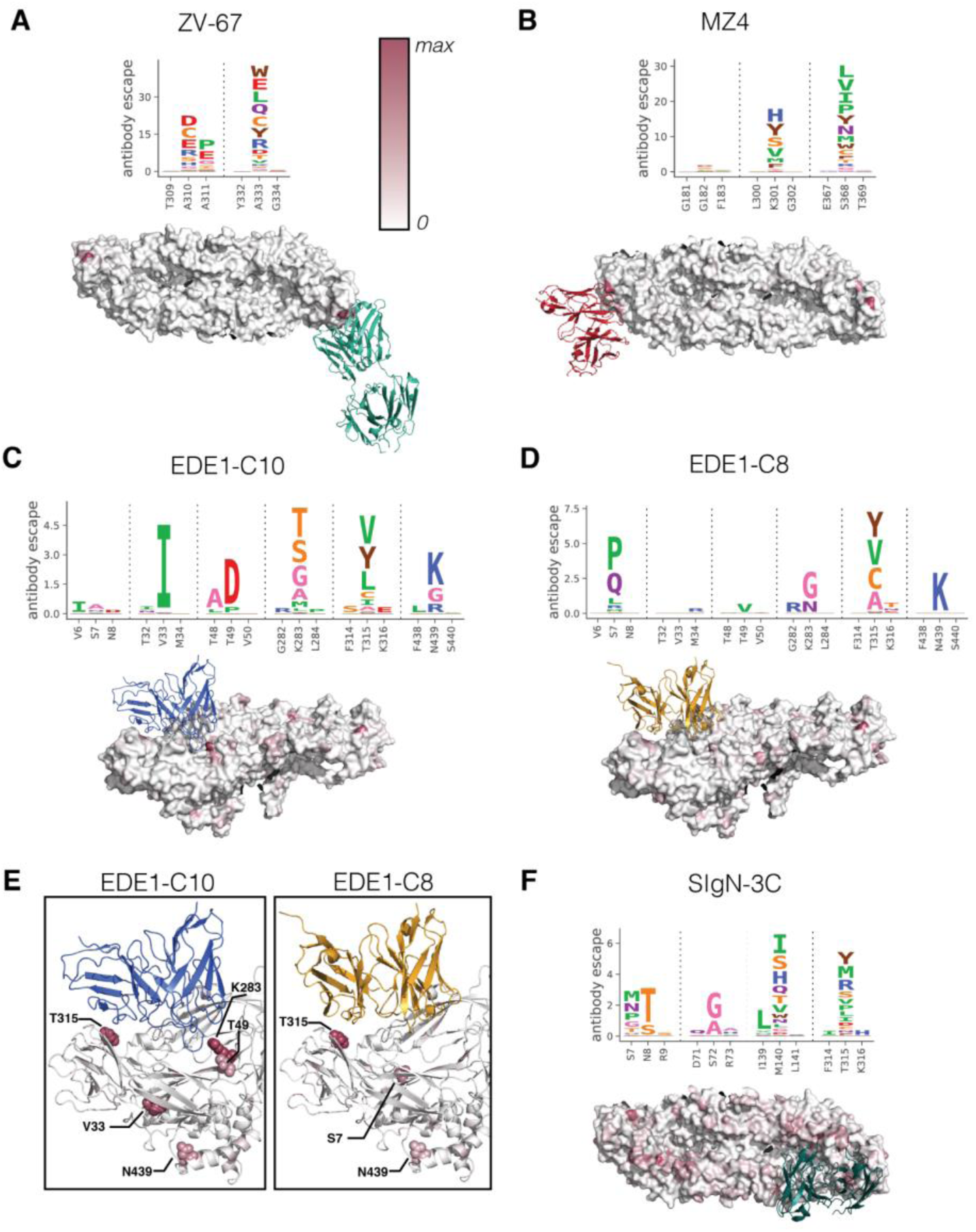
Site and mutational-level antibody escape visualized as logo plots and projected on protein structures. In panels **A-D** and **F**, the mutation-level antibody escape is shown as logo plots where the height of the letter is scaled to the magnitude of antibody escape attributed to that amino acid mutation. Below the logo plots, the site-level antibody escape values are used to color the E protein structure from white to red, and are scaled per antibody. **(E)** Zoomed-in sites of escape for class EDE1 antibodies.

In contrast, broadly neutralizing antibodies were escaped by fewer mutations of weaker effects both within and outside structural epitopes. Despite similar structural epitopes^24–26,28^, EDE1-C10 and EDE1-C8, were escaped by mutations at distinct sets of residues. The EDE1-C10 functional epitope involved the apparent antibody binding footprint (sites T49, R283, and T315)^24^, as well as a site outside the binding footprint at the intra-dimer interface (site V33) (**Figure 3C**). EDE1-C8 was escaped by distinct mutations near the binding footprint and at the intra-dimer interface, with the highest degree of escape occurring at sites S7 and T315 (**Figure 3D**). Interestingly, S7P, one of the mutations with the largest effect for EDE1-C8, and which conceivably introduces a major structural change with a proline substitution in the homodimer core, did not cause appreciable escape from EDE1–C10 (**Figure C,D**). In fact, there is very little overlap in the sets of escape sites from these two antibodies, despite substantial overlap in their structural epitopes (**Figure 3C-E**).

Another broad antibody, SIgN-3C, had escape mutations distributed across E protein domains. We identified sites S72, M140 and T315 as those comprising the functional epitope, where few mutations had large individual effects (**Figure 3F**). These functional epitope residues fell within or nearby the previously described structural epitope^23^. Notably, Zhang et al. showed that SIgN-3C Fab is complexed with E homodimers in 3 distinct conformations, thus widely distributing interaction requirements across E protein^23^. Our data also shows signs of lower level selection across other components of these 3 variable epitopes; the bc loop (82-83), the kl loop (279-281) and nearby residues 315-316 at the intra-dimer interface. This widely distributed and flexible epitope could be the cause of the relatively high levels of noise in this dataset (**Supplemental Figure 6**), particularly when compared with antibodies with simpler epitopes (**Supplemental Figures 2, 3**).

### Validations in single mutant neutralization assays

To validate the findings from our deep mutational scanning, we made single mutations in Zika virus E protein and tested them in dose-response neutralization assays (**Figure 4**). These individual assays allowed us to examine antigenic effect size across a range of antibody concentrations, rather than a single, potent concentration. As we sought to measure the effects of many mutations in high-throughput, we validated mutations in a previously described pseudovirus system that maintains antigenicity of flavivirus particles^1,36^.

**Figure 4.**
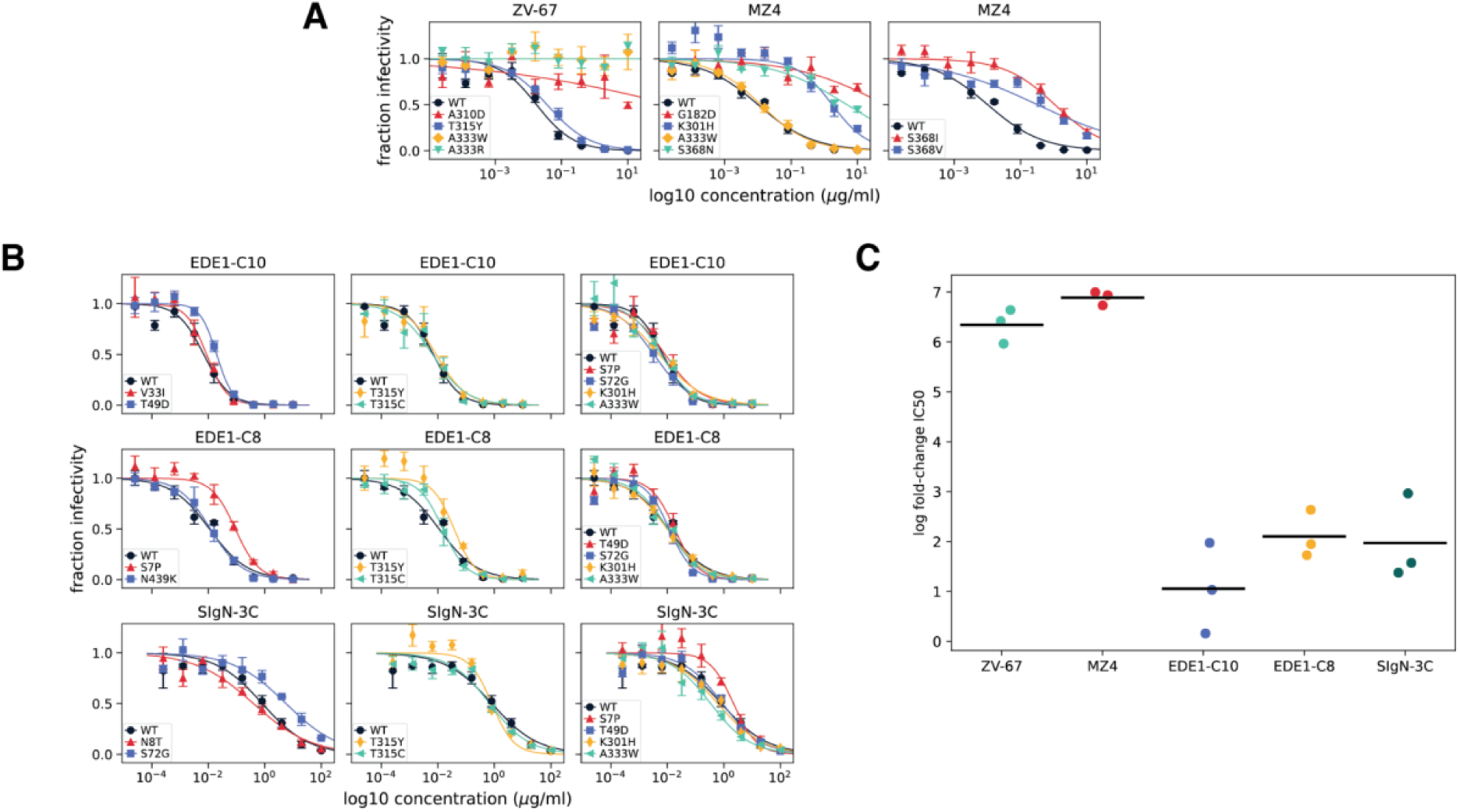
Validation of single E mutations in historic Zika virus strain MR766. Representative neutralization assays demonstrating large effects of mutations against narrow and pseudo-broad antibodies, and small effects against broad antibodies **(A)** Narrow and pseudo-broad neutralizing antibodies tested in dose-response neutralization assays against Zika virus MR766 mutants. The mutations A310D, A333W and A333R are expected to escape ZV-67, whereas T315Y should not affect neutralization. Against MZ4, the mutations G182D, K301H, S368N, S368I, and S368V are all expected to escape, while A333W should have no effect. **(B)** Dose-response neutralization assays of viral mutants against broad antibodies. The first panel contains mutations expected to uniquely escape each antibody: V33I and T49D for EDE1-C10; S7P and N439K for EDE1-C8; and N8T and S72G for SIgN-3C. The second panel contains mutations T315Y and T315C. While T315Y is expected to modestly increase neutralization resistance to all three broadly neutralizing antibodies, only T315C is expected to modestly increase resistance specifically to EDE1-C8. The third column contains mutations not expected to increase neutralization resistance against the indicated antibody. In all panels, points indicate the mean and standard error across three technical replicates. **(C)** Comparing the log fold-change in IC50 of mutant viruses with the single largest effect against each antibody, relative to wild-type. Points refer to single log fold-change neutralization assay replicates and lines indicate the mean of those replicates.

We first assessed the effects of mutations identified in deep mutational scanning of narrow and pseudo-broad antibodies ZV-67 and MZ4 (**Figure 4A**). We saw that mutations identified by ZV-67 selection completely escaped neutralization (**Figure 4A; Supplemental Table 1**). For MZ4, we validated several escape mutations identified by our deep mutational scanning. A small subset of these mutations (G182D and S368N) were previously identified by passaging experiments with a contemporary ‘Asian’ lineage strain of Zika virus (Paraiba_01), different from the ‘African’ lineage strain used as the basis of our libraries (MR766). Deep mutational scanning identified both the mutation S368N as well as other S368 mutations and K301H as potential escape mutations, all of which validated with large effects in neutralization assays (**Figure 4A; Supplemental Table 3**). While our traditional neutralization assays validated the effect of G182D in Zika pseudovirus strain MR766, deep mutational scanning of MZ4 did not identify the large effect size of this mutation. Together, we observed that single mutations had large effects on ZV-67 and MZ4 neutralization, either completely ablating neutralization or significantly increasing neutralization resistance.

For broadly neutralizing antibodies, single mutations had far more modest effects. As above, we generated mutant Zika virus MR766 pseudovirus particles and measured their effects in neutralization assays. Antibody selection with EDE1-C10 had identified the mutations V33I at the intra-dimer interface and T49D within the structural epitope (**Figures 3C,E**). Our neutralization assays demonstrated T49D modestly but significantly affected neutralization resistance, while the effects of V33I on neutralization resistance were minimal or undetectable (**Figure 4B**; **Supplemental Table 4**). We also tested the mutation T315Y, which had a smaller mutation-level antibody escape value, but occurred within a site of larger summed antibody escape. We saw T315Y behaved similarly to T315C, a negative control mutation expected to have no effect on neutralization. Finally, we also tested EDE1-C10 against a variety of mutations identified in deep mutational scanning with other antibodies in our panel, and saw no significant effects on neutralization.

Deep mutational scanning of another broad antibody, EDE1-C8, selected mutations S7P and N439K outside its structural epitope (**Figures 3D,E**). When we tested these mutations in dose-response neutralization assays against EDE1-C8, we saw that S7P significantly increased neutralization resistance, whereas N439K was neutralized similarly to wild-type (**Figure 4B; Supplemental Table 5**). Again, we tested mutations T315Y and T315C, which had lower mutation-level antibody escape from EDE1-C8 (**Figure 3D**). We saw small magnitude, non-significant effects on neutralization. As expected, when we tested EDE1-C8 against a panel of negative control mutations, we saw minimal effects on neutralization.

Finally, we tested the broad antibody SIgN-3C against a panel of predicted escape mutations, N8T, S72G and T315Y. To a certain extent, the effect size of these mutations identified from our deep mutational scanning data (**Figure 3F**) correlated with the effect size quantified by traditional neutralization assay. For example, the mutation S72G significantly increased neutralization resistance, whereas T315Y, which had a much smaller antibody escape value was indistinguishable from wild-type (**Figure 4B; Supplemental Table 6**). However, the mutation N8T, which had a larger antibody escape value, was also neutralized similarly to wild-type.

Overall, these neutralization assays validate the finding from the deep mutational scanning that narrow and pseudo-broad antibodies have very large-magnitude escape mutations, while no mutations more than modestly affect neutralization by broad antibodies (**Figure 4C**). For the narrow antibodies, the change in neutralization caused by specific mutations was well predicted by the deep mutational scanning For the broad antibodies, there was often correspondence between the deep mutational scanning and validation neutralization assays, but the mutation effects were small in both assays, and in some cases undetectable in the neutralization assays.

### Mutational tolerance at functional epitopes of broad and narrow neutralizing antibodies

Together, our deep mutational scanning and traditional neutralization assays showed many single mutations across a few sites had large effects on neutralization by antibodies with narrow specificities. However, for broadly neutralizing antibodies, we identified only a few mutations across many E protein sites with small effects. Was this because broad antibodies tended to be escaped by mutations that were not well tolerated in the replicative viruses used in our libraries? Or because broadly neutralizing antibodies distribute interaction requirements over a wide range of sites?

To address our first question, we leveraged data published in Sourisseau et al^34^ assessing the effects of all MR766 E protein mutations on virus infectivity *in vitro.* We compared these mutational tolerance values against mutation-level antibody escape in **Figure 5A**. Interestingly, we saw that mutations selected by broad antibodies were not uniformly poorly tolerated. In fact, a few mutations that escaped EDE1-C8 were relatively well tolerated.

**Figure 5.**
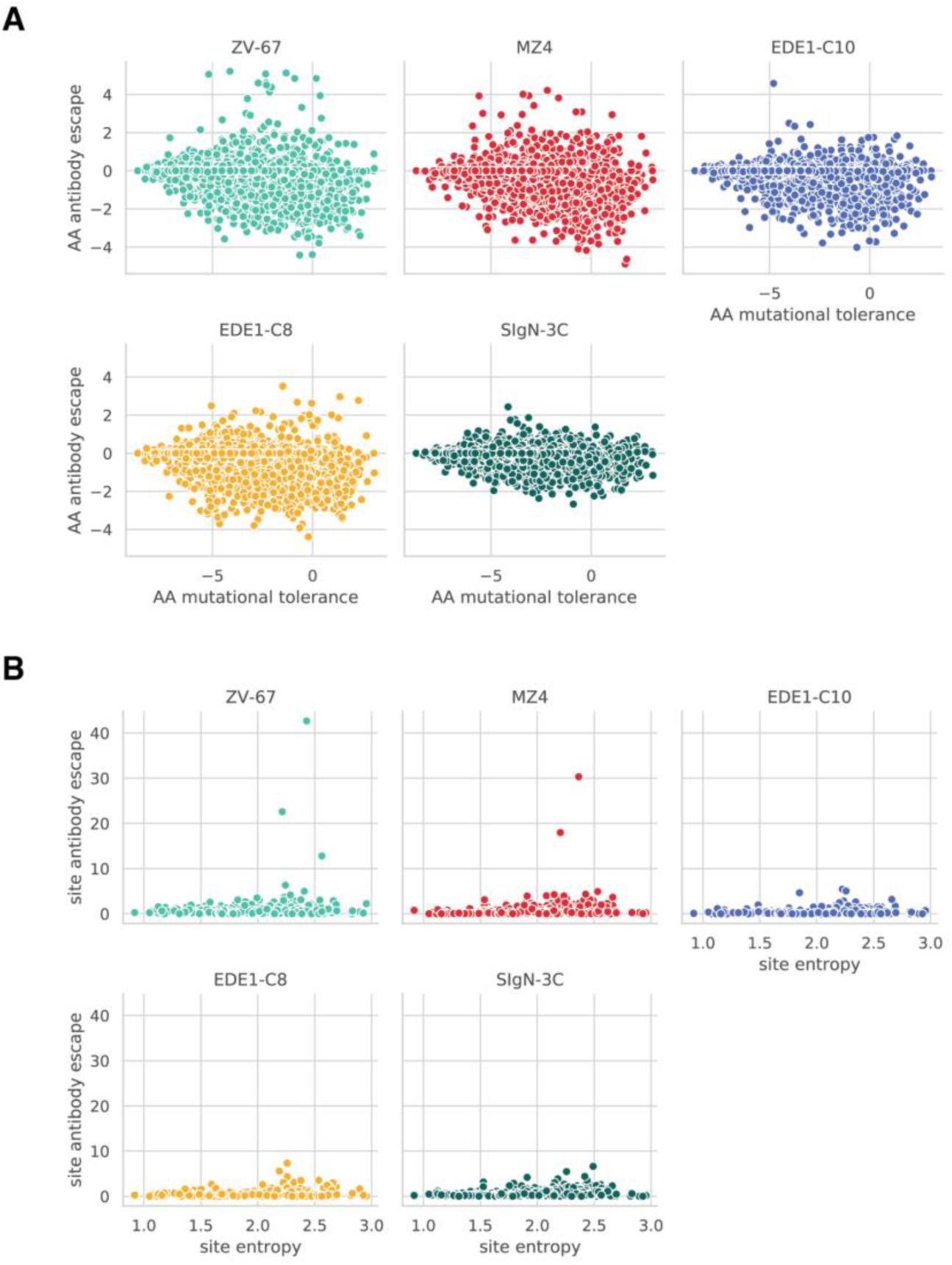
Mutational tolerance at the mutation- and site-level for broad and narrow neutralizing antibodies. **(A)** Amino acid (here abbreviated as AA) antibody escape plotted against amino acid mutational tolerance. Mutational tolerance was calculated by taking the log ratio of mutant amino acid preference over wild-type amino acid preference from a previously published deep mutational scanning study^34^. A mutational tolerance score of 0 indicates functionality similar to the wild-type amino acid at that site, while positive and negative mutational tolerance values indicate increased and decreased functionality, respectively. See **Methods** for more details. **(B)** Site-level escape values for all antibodies plotted against site-level entropy. For a similar plot showing a different site-level mutational tolerance metric, see **Supplemental Figure 8.**

Crucially, despite being well tolerated, these mutations still conferred relatively low degrees of antibody escape. This becomes particularly evident when compared to antibodies with narrow specificities, like ZV-67, where escape mutations of variable tolerance consistently conferred high degrees of antibody escape.

Next, we examined site-level data to see if interaction requirements were distributed differently between antibodies with broad and narrow specificities. To estimate site-level mutational tolerance, we used two metrics: number of effective amino acids and site entropy. These values were plotted against site-wise summed antibody escape values for all antibodies in **Figure 5B**. These data show that for antibodies with narrower specificities, i.e. MZ4 and ZV-67, antibody escape is focused at a few relatively mutationally tolerant sites. Broadly neutralizing antibodies EDE1-C10, −C8 and SIgN-3C exhibit distributed antibody escape across sites of varying mutational tolerance. Altogether, these data support both proposed explanations for a lack of large-effect single amino acid escape mutations to broadly neutralizing antibodies. Broadly neutralizing antibodies target both constrained and mutationally tolerant sites, but in all cases are only weakly escaped by a few mutations at a given site.

### Antigenic effect of escape mutations in other flavivirus genetic contexts

We next tested if the antigenic effects of mutations we had identified and validated in Zika virus MR766 were similar in other flaviviruses. Zika virus MR766 is closely related to a widely-utilized contemporary strain of Zika virus, H/PF/2013, sharing ∼96% amino acid E sequence identity. It is more distantly related to dengue viruses, sharing only ∼54% identity with dengue virus serotype 2 (strain 16681). Nevertheless, as the broad and pseudo-broad antibodies within our panel neutralized this wide variety of viruses, we hypothesized some degree of antigenicity might be conserved.

To test this hypothesis, we generated single mutant pseudovirus particles of Zika virus strain H/PF/2013 and dengue virus serotype 2 strain 16681. For each antibody, we selected a single viral E mutation that escaped Zika virus strain MR766 (as validated in **Figure 4**). We then examined a multisequence protein alignment (**Figure 6A**) for the corresponding residues in Zika virus strain H/PF/2013 and dengue virus serotype 2 strain 16681. Where sequence was not conserved, we still mutagenized to the predicted escape residue from our deep mutational scanning. Because this wild-type sequence is not conserved, in **Figure 6B** we only refer to mutations by site and mutant amino acid code (e.g., ‘S7P’ is now abbreviated to ‘7P’). Additionally, we accounted for the small deletions in dengue virus E relative to Zika virus, ensuring the same antigenic regions were mutagenized.

**Figure 6.**
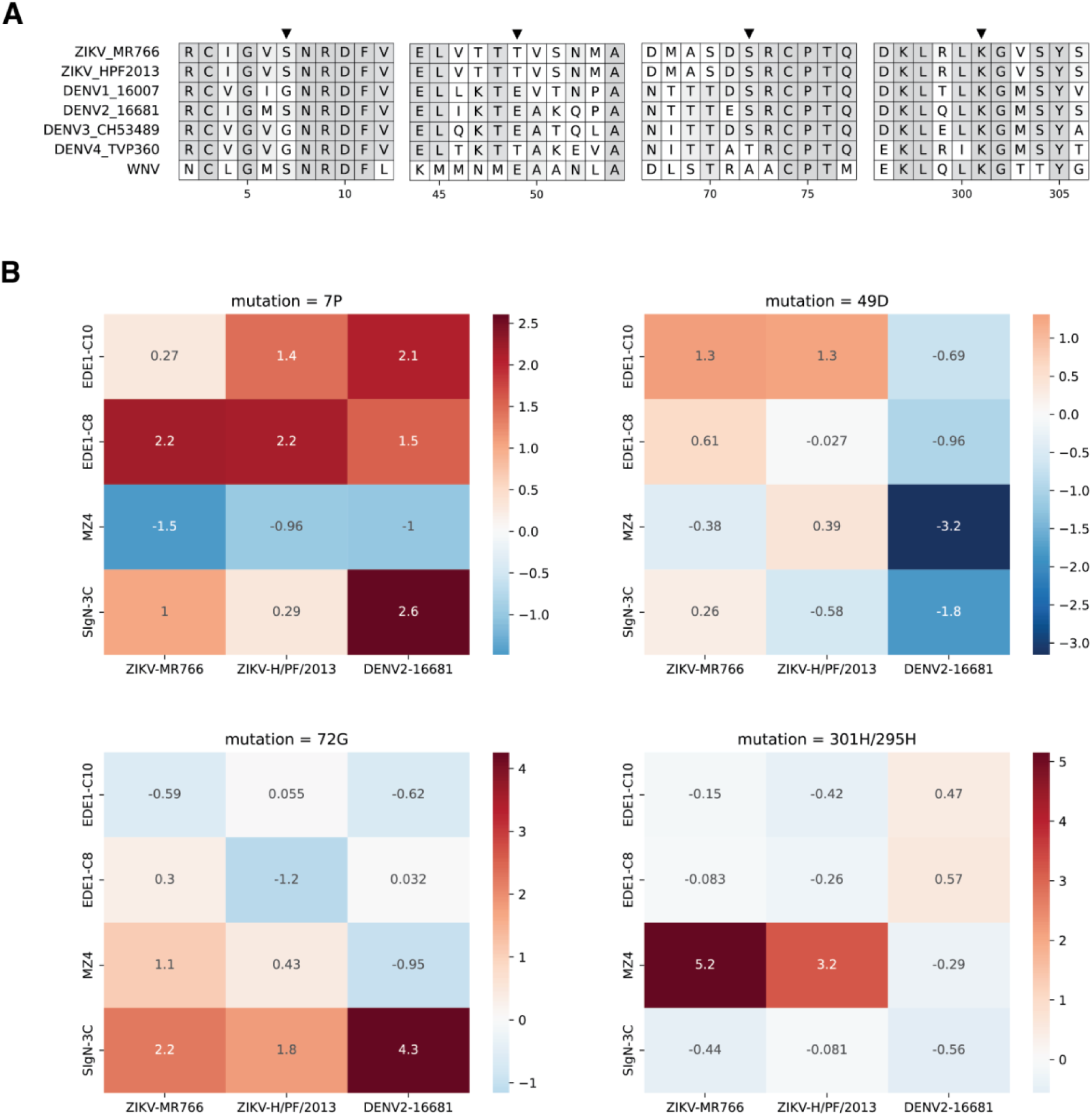
Validation of single E mutations in other flavivirus genetic contexts. **(A)** A subset of amino acid alignment of Zika virus (strain MR766), Zika virus (strain H/PF/2013), dengue virus serotype 1 (strain 16007), dengue virus serotype 2 (strain 16681), dengue virus serotype 3 (strain CH53489), dengue virus serotype 4 (strain TVP360), and West Nile virus. Gaps in sequence reflect omitted sections of protein alignment. For full alignment, see **Supplemental Figure 7.** Black arrows indicate sites where site-directed mutagenesis was targeted. The mutation 7P is expected to escape EDE1-C8, 49D is expected to escape EDE1-C10, 72G is expected to escape SIgN-3C, and 301H is expected to escape MZ4. Light gray highlight indicates site percent identity. **(B)** Dose-response neutralization assays against Zika virus (strain MR766), Zika virus (strain H/PF/2013) and dengue virus serotype 2 (strain 16681) reporter pseudovirus engineered with single amino acid mutations predicted to confer escape in Zika virus (strain MR766). Heatmaps colored red to blue by the log fold-change in IC50 between a given viral mutation and wild-type, and scaled per viral mutation. Larger positive values (red) indicate increased neutralization resistance or viral escape, whereas negative values (blue) indicate neutralization sensitization. For the mutation 301H, a deletion in dengue virus relative to Zika virus adjusts this mutation identity to 295H. The corresponding neutralization assays summarized in this figure are shown in **Supplementary Figures 8, 10 and 11**.

Next, we again performed dose-response neutralization assays and calculated the log fold-change from wild-type to mutant IC50s. These values are shown as heatmaps in **Figure 6B**. For EDE1-C8 and EDE1-C10, deep mutational scanning identified the domain I escape mutations 7P and 49D, respectively. Across flavivirus backgrounds, the mutation 7P confers minute-to-modest increases in neutralization resistance against all broadly neutralizing antibodies. Against the pseudo-broadly neutralizing antibody MZ4, 7P is modestly sensitizing. In contrast, 49D only increased resistance to EDE1-C10 neutralization in Zika virus strains, but not in dengue virus. As previously mentioned, the mutation 7P exists at the intra-dimer interface, whereas 49D falls within the binding footprint of EDE1-C10. This potentially explains the relatively conserved effect of 7P relative to 49D, as introducing a proline residue at a critical protein-protein interface might have greater antigenic effects than single charge alterations within a binding footprint. Similarly to EDE1-C10, for MZ4, we saw that the effect of its identified escape mutation K301H was only conserved within Zika virus (**Figure 6B**)

Against SIgN-3C, the mutation 72G conferred neutralization resistance from both strains of Zika virus and dengue virus serotype 2 (**Figure 6B**). Previously, Zhang et al. reported that SIgN-3C neutralizes Zika virus and dengue viruses by different mechanisms: either by aggregating virus particles (Zika) or inhibiting endosomal fusion (dengue)^23^. In combination with their structural studies, our findings suggest that the contribution of 72G is independent of the neutralization mechanism. In summary, these data indicate potential conservation of functional epitopes across divergent flaviviruses of some – but not all – broadly neutralizing antibodies.

## Discussion

Here, we used deep mutational scanning to quantify the effect of all possible amino acid mutations in Zika virus E protein on neutralization by antibodies with broad and narrow specificities. Our results indicate that narrow antibodies are quantifiably easier to escape than broad antibodies. Specifically, there are single amino-acid mutations that can completely ablate neutralization by narrow strain-specific antibodies, whereas no mutation more than modestly reduces neutralization by the broad antibodies. This finding is reminiscent of prior work comparing the ease of escape from broad and narrow influenza neutralizing antibodies, where the identified mutations had small effects against broad antibodies relative to large effects against narrow antibodies^35^. We also saw that the escape mutations we identified, which comprise antibody functional epitopes, did not completely overlap amongst antibodies with highly similar structural epitopes. Thus, our work further provides evidence of multiple highly conserved broadly neutralizing functional epitopes.

Furthermore, for broadly neutralizing antibodies, we show that functional epitopes can be composed of residues outside of the structural epitope. One explanation for this finding could be that mutations at the intra-dimer interface destabilize the E homodimer interaction required for broadly neutralizing antibody epitopes, which are typically quaternary. This explanation would be consistent with evidence suggesting that EDE1 class antibodies neutralize Zika and dengue virus by stabilizing E proteins as prefusion homodimers^24,28^. This subsequently prevents the conformational changes necessary for E protein to initiate fusion with host cell membranes^37^. Additionally, our data also indicate that the antigenic effects of escape mutations are conserved across an array of diverged flavivirus E proteins, concordant with a model of antibody interaction at conserved residues across divergent antigens.

Previous efforts to characterize antigenicity of flavivirus surface proteins have not been able to distinguish between functionally required residues for neutralization versus binding^23–26,28^. Other efforts, such as phage display platforms to perform alanine scanning of dengue virus and small scale site-directed mutagenesis could only identify linear antibody epitopes, and only provide limited information on a few domains of E^16,17,38^. In contrast, our approach comprehensively measures the effect of all single amino acid mutations on neutralization by antibodies with complex quaternary epitopes. Any mutations outside of structural epitopes that affect antigenicity and reduce neutralization – such as those identified herein – are key considerations for vaccine design utilizing broadly neutralizing antibodies as prototypes.

While our work examines the most accessible form of viral genetic diversity – single amino acid mutations – whether these mutations act in combination with each other in any viral genetic context remains to be seen. Future work examining epistatic effects of mutations will be key to understanding the potentially compounding effects of any of the effects of mutations measured here. Another caveat of our findings was the level of noise observed in deep mutational scanning of the broad antibodies EDE1-C10 and SIgN-3C. Previously, structural studies have indicated these antibodies bind flavivirus E protein in three slightly different conformations, potentially creating a pseudo-polyclonal response^23,26^. This more distributed selective force could lead to the wide-ranging and more noisy antibody escape in both SIgN-3C- and EDE1-C10-selected libraries. For EDE1-C8, no structural data has produced similar findings, perhaps explaining why the deep mutational scanning produced less noise.

The deep mutational scanning reported here will be of benefit towards a number of applications. Broadly flavivirus-neutralizing antibodies have been protective against lethal disease and reduced viral load in murine models^32,39,40^, indicating as-yet untapped therapeutic potential of these proteins. With the effects of global warming and urbanization widening the endemic range of the Zika and dengue virus mosquito vectors and increasing overlap with sylvatic transmission cycles, epidemic preparation is paramount^41–43^. If the broadly neutralizing antibodies tested here were easily escaped, the attention and depth of characterization they receive would be difficult to rationalize. Our work therefore highlights the utility of broadly neutralizing antibodies as protective agents, but also stresses the importance of examining the potential of these new therapies to be escaped by viral mutations. Pertinent to public health efforts monitoring new and emerging flavivirus variants, a few escape mutations described here have relatively conserved antigenic effects across divergent flaviviruses. Future development of deep mutational scanning libraries in other strains of Zika and dengue viruses could more deeply characterize how conserved these effects may be.

## Acknowledgements

We thank John Huddleston for the helpful discussion and advice; Robin J. Kaai for technical assistance; Theodore C. Pierson for providing Raji/DC-SIGNR cells and plasmids for reporter virus particle production; Michael S. Diamond for providing the antibody ZV-67; and Shelly Krebs for providing expression plasmids for the antibody MZ4.

This work was supported by the Viral Pathogenesis and Evolution Training Grant T32 AI083203 to CK; the Diseases of Public Health Importance Training Grant T32 AI007509 to JBS; the Fred Hutchinson Cancer Center Diverse Trainee Fund to MC; the Fred Hutch Scientific Computing (NIH grants S10-OD-020069 and S10-OD-028685); the Fred Hutch Genomics & Bioinformatics Shared Resource (RRID:SCR_022606); the Fred Hutch Antibody Technology Core (RRID:SCR_022608); and the Shared Resource Facilities of the Fred Hutch/University of Washington Seattle Children’s Cancer Consortium (P30 CA015704). JDB is an Investigator at the Howard Hughes Medical Institute.

## Author contributions

JDB and LG conceived of the study, and ME helped develop the Zika virus mutant libraries. CK, CHCA, JBS, MC and LL and performed the experiments. CK wrote the first manuscript draft. All authors reviewed and provided comments on the manuscript.

## Supplementary Information

**Supplemental figure 1.**
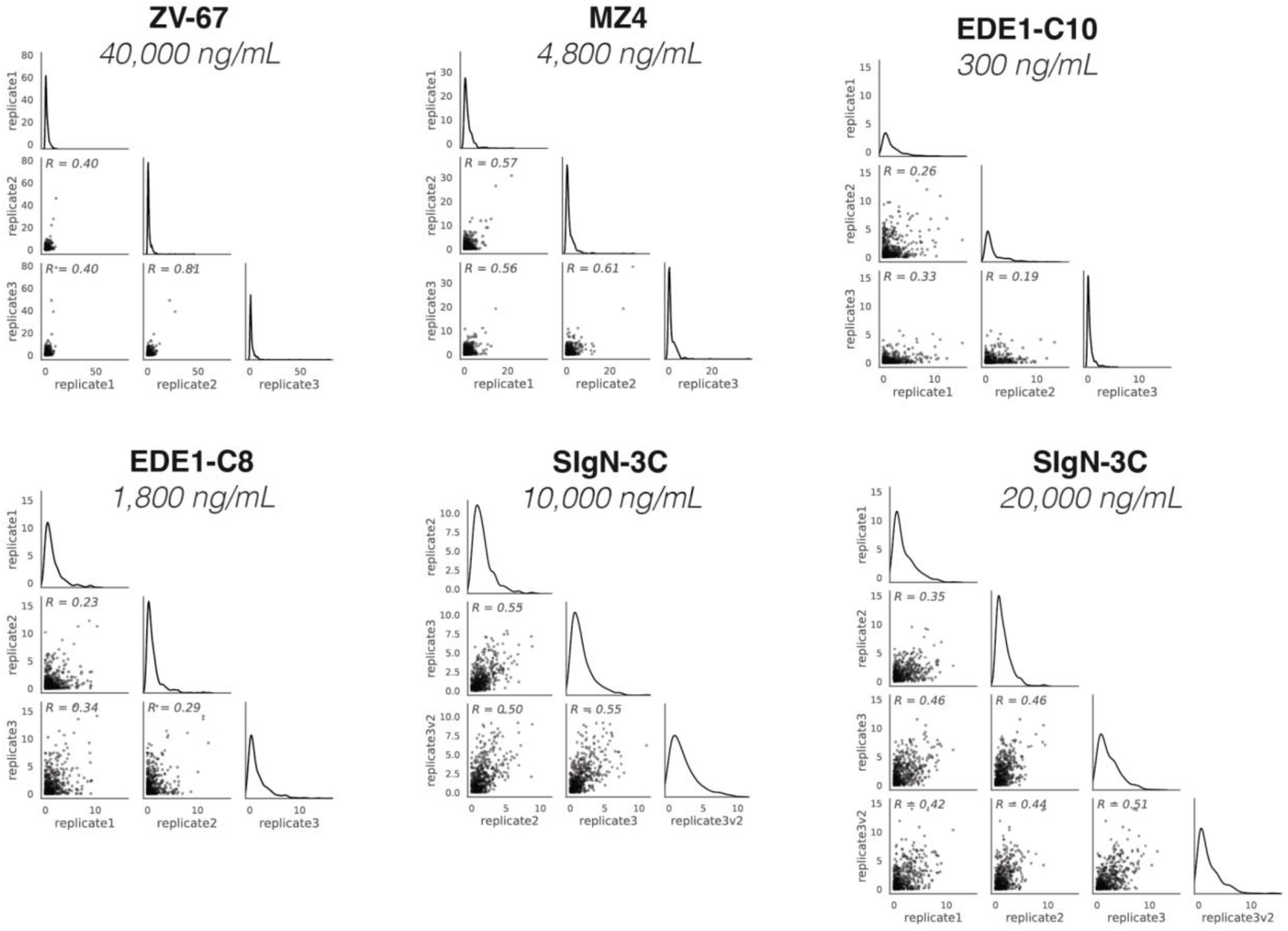
Correlations across replicate deep mutational scanning experiments. Points represent site-wise differential selection (our metric of antibody escape) across all amino-acid mutations. The replicates correlate well for narrow antibodies, where a few sites lead to high degrees of escape. For broad antibodies, where no sites lead to large magnitudes of escape, the results are reasonably correlated.

**Supplemental figure 2.**
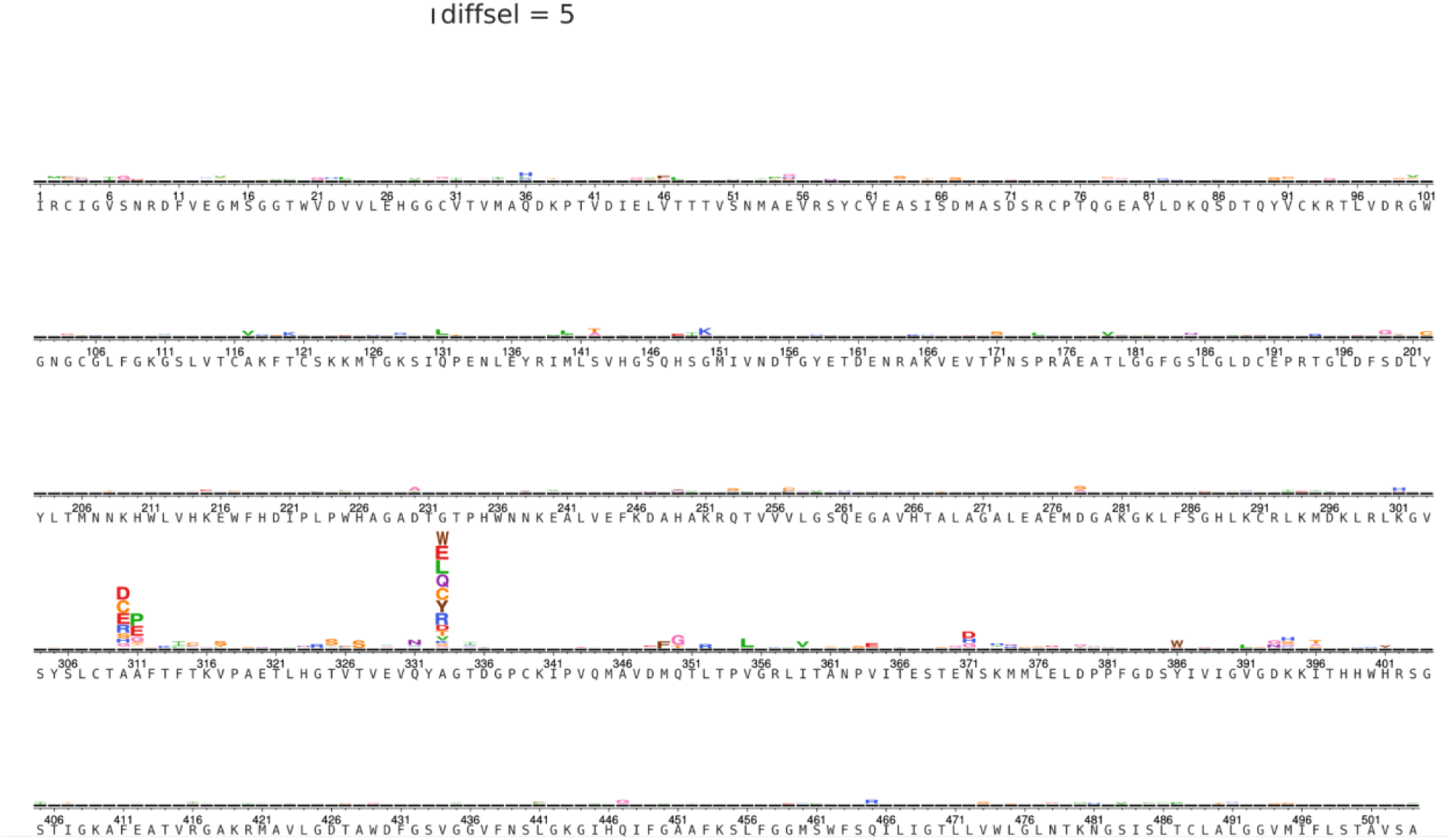
Mutation-level deep mutational scanning from ZV-67 selections across the entire E protein. The effect of single amino acid mutations on neutralization, shown as logo plots where the height of the letter is scaled to the magnitude of antibody escape attributed to that amino acid mutation. The plot is scaled by differential selection (diffsel), our metric of antibody escape. See **Methods** for more details.

**Supplemental figure 3.**
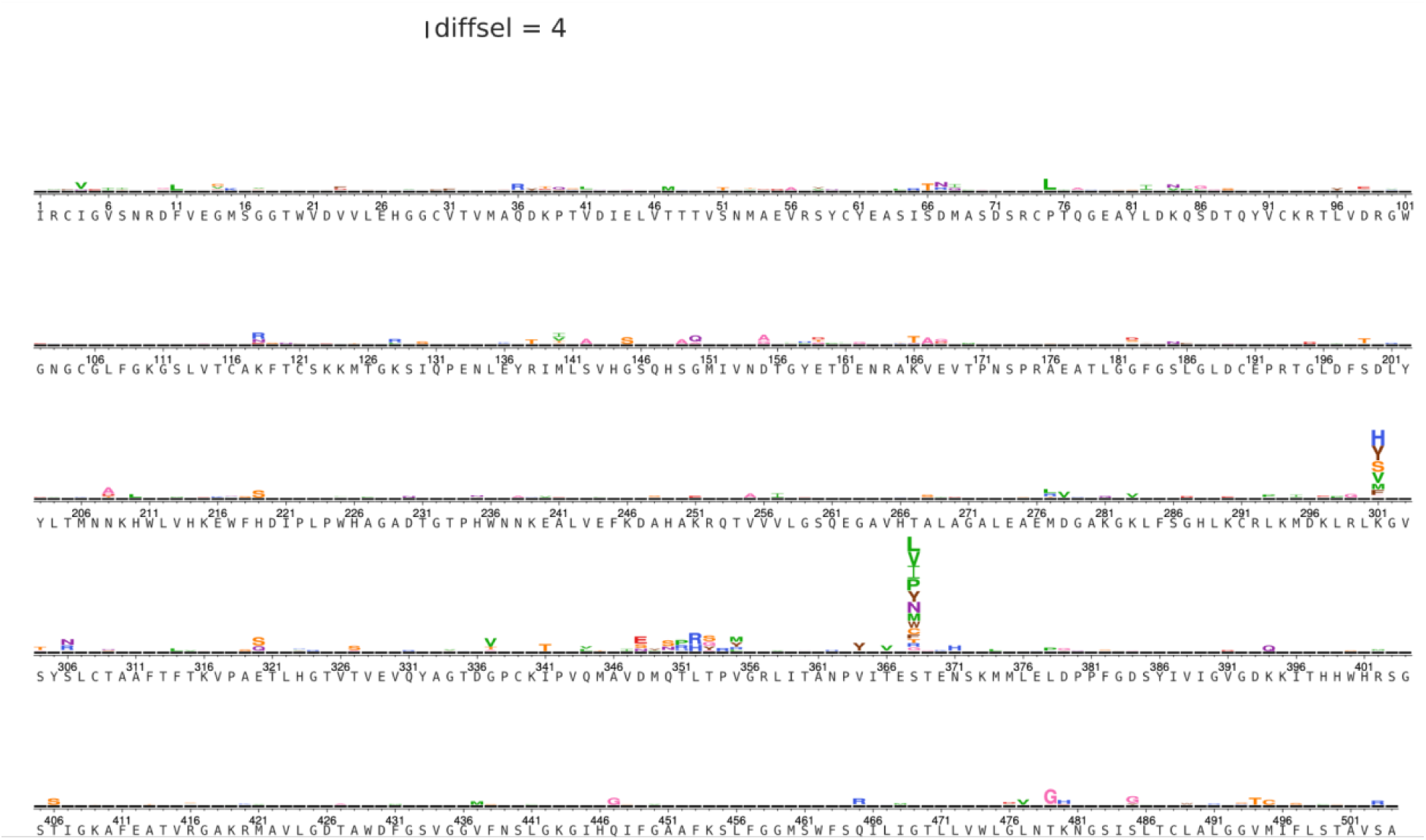
Mutation-level deep mutational scanning from MZ4 selections across the entire E protein. The effect of single amino acid mutations on neutralization, shown as logo plots where the height of the letter is scaled to the magnitude of antibody escape attributed to that amino acid mutation. The plot is scaled by differential selection (diffsel), our metric of antibody escape. See **Methods** for more details.

**Supplemental figure 4.**
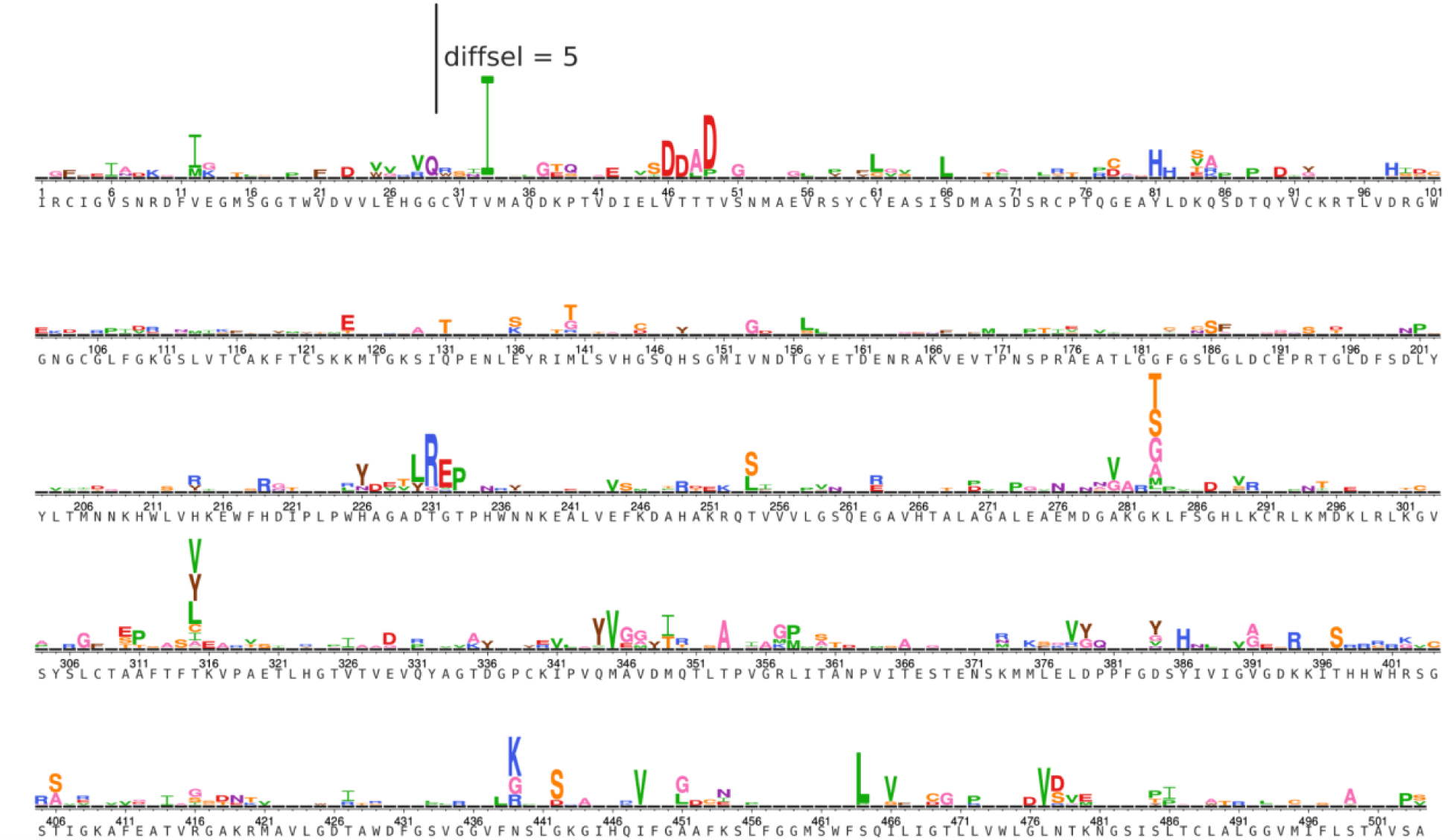
Mutation-level deep mutational scanning from EDE1-C10 selections across the entire E protein. The effect of single amino acid mutations on neutralization, shown as logo plots where the height of the letter is scaled to the magnitude of antibody escape attributed to that amino acid mutation. The plot is scaled by differential selection (diffsel), our metric of antibody escape. See **Methods** for more details.

**Supplemental figure 5.**
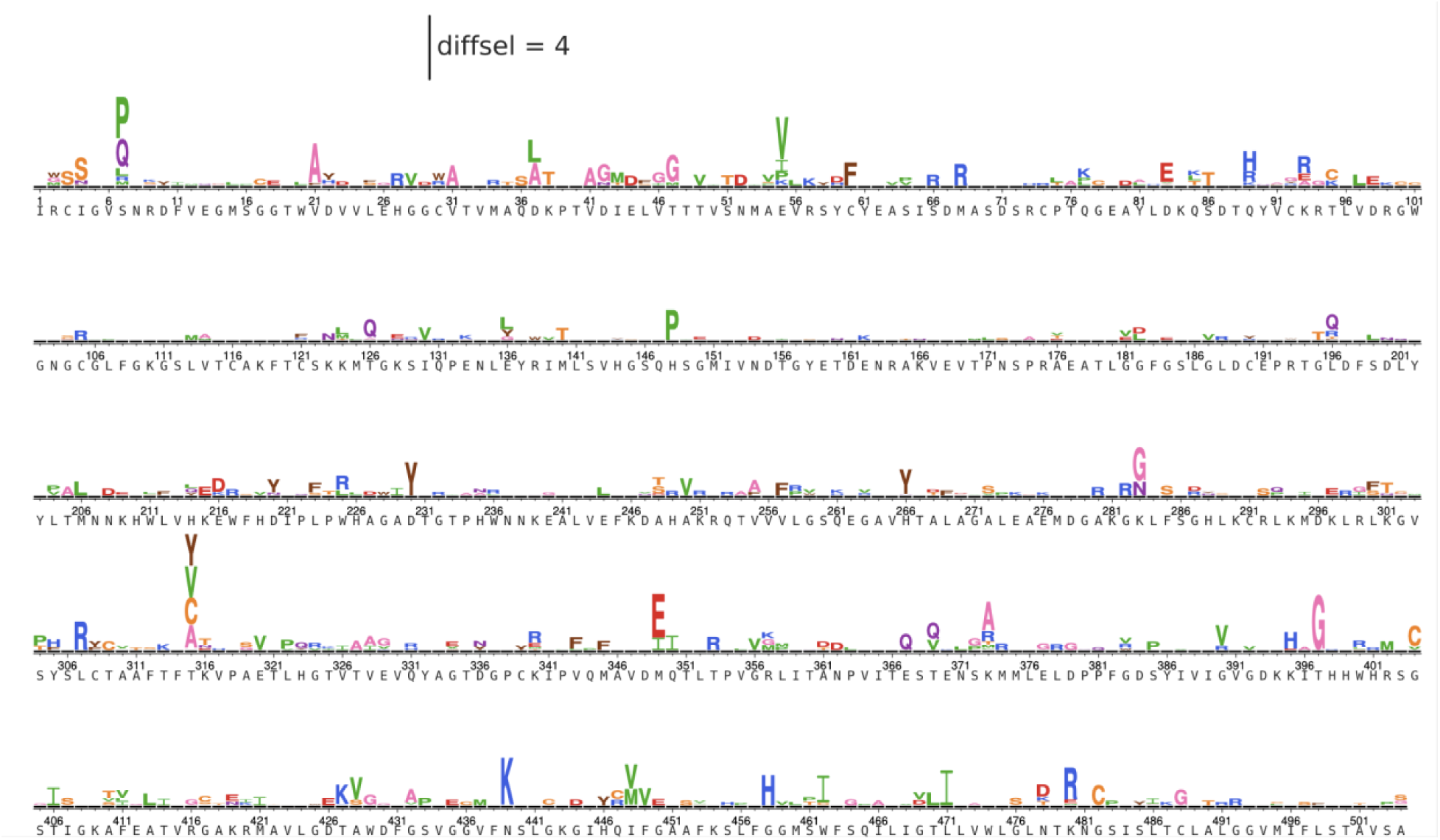
Mutation-level deep mutational scanning from EDE1-C8 selections across the entire E protein. The effect of single amino acid mutations on neutralization, shown as logo plots where the height of the letter is scaled to the magnitude of antibody escape attributed to that amino acid mutation. The plot is scaled by differential selection (diffsel), our metric of antibody escape. See **Methods** for more details.

**Supplemental figure 6.**
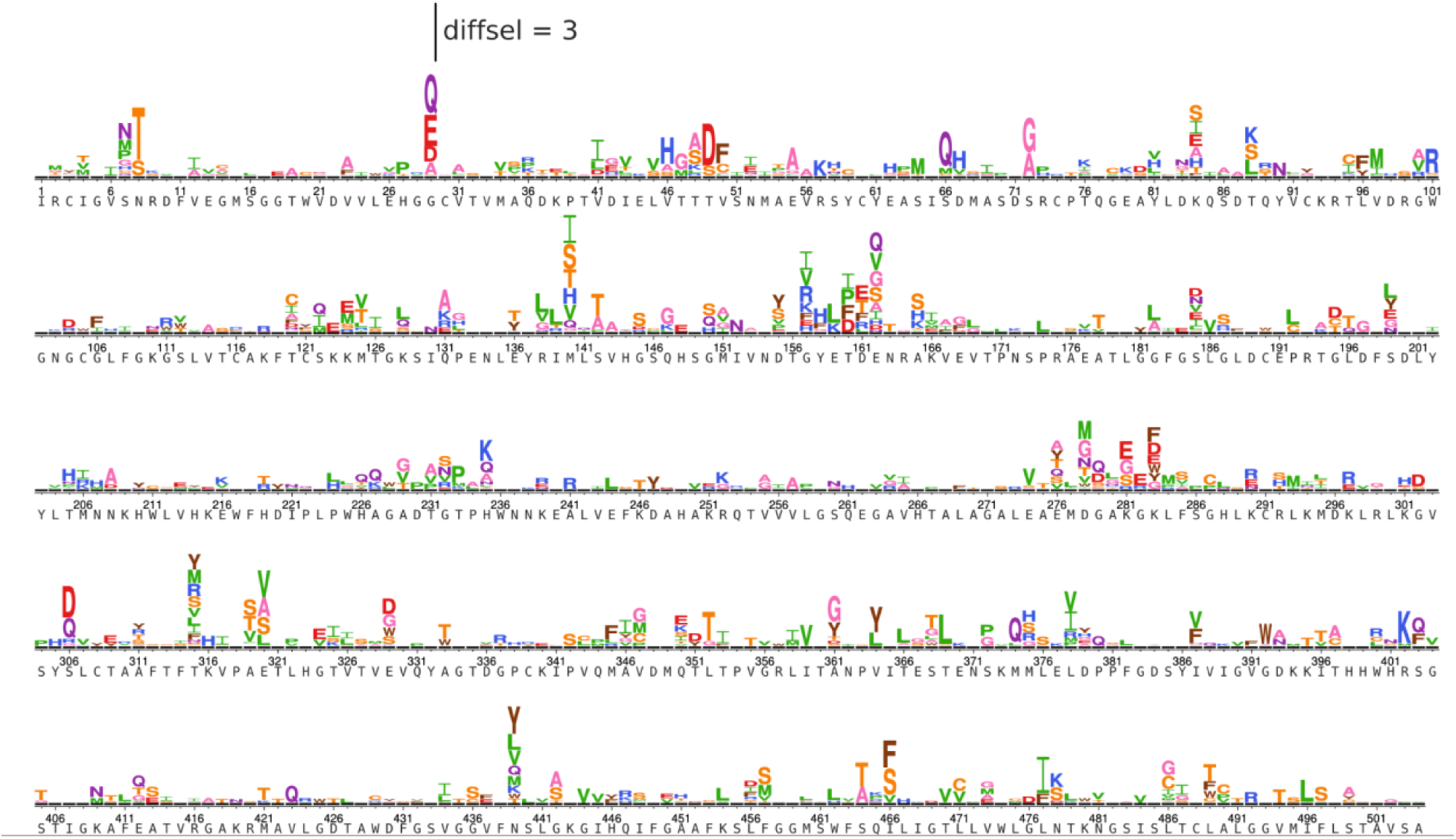
Mutation-level deep mutational scanning from SIgN-3C selections across the entire E protein. The effect of single amino acid mutations on neutralization, shown as logo plots where the height of the letter is scaled to the magnitude of antibody escape attributed to that amino acid mutation. The plot is scaled by differential selection (diffsel), our metric of antibody escape. See **Methods** for more details.

**Supplemental figure 7.**
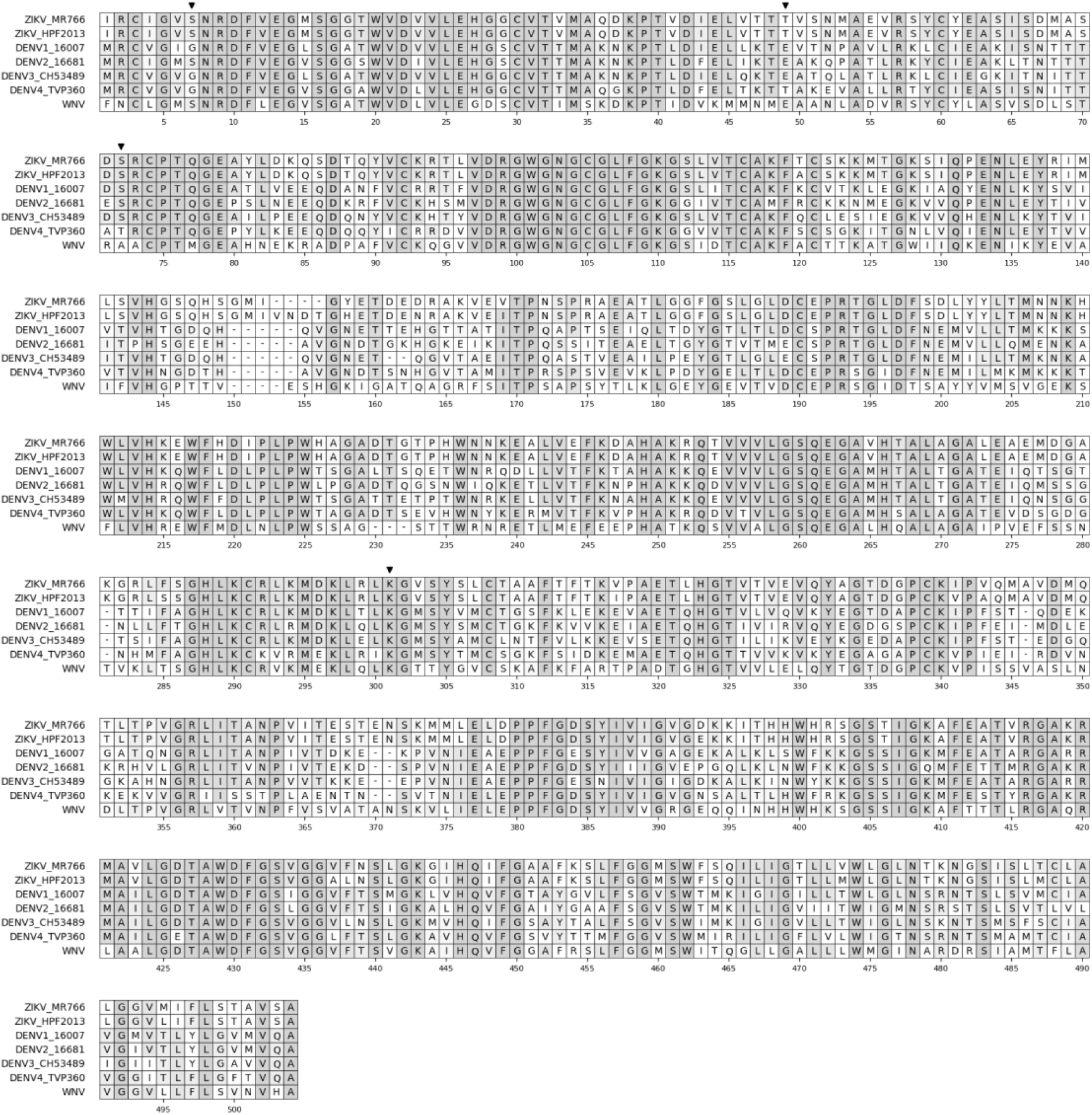
Flavivirus E protein alignment. Black arrows indicate sites where site-directed mutagenesis was targeted, and light gray highlight indicates site percent identity. See **Methods** for more details on sequences used and how the alignment was generated.

**Supplemental figure 8.**
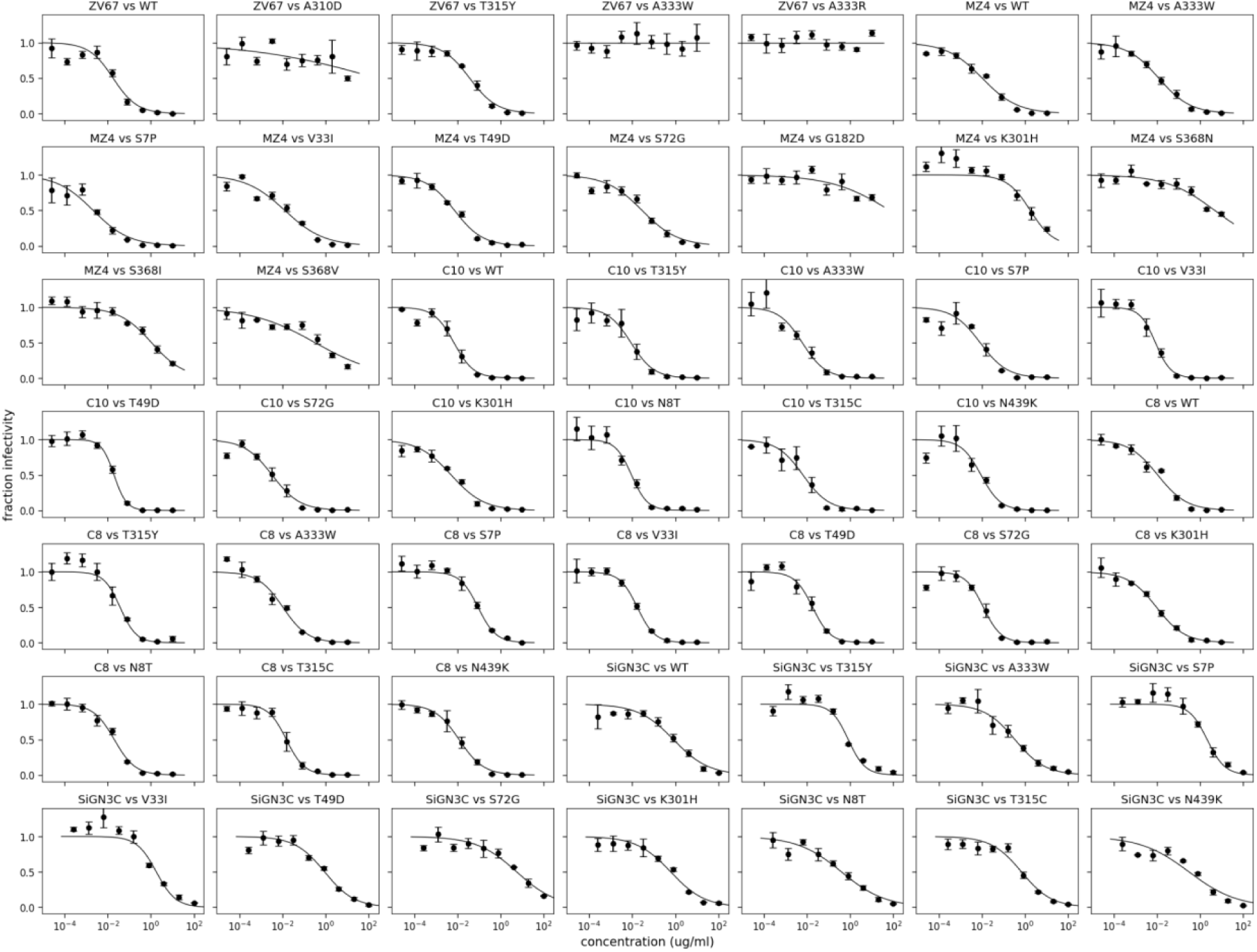
Individual Hill Curves interpolated for technical triplicate neutralization assays with Zika virus MR766. Points indicate the mean and standard error across three technical replicates.

**Supplemental figure 9.**
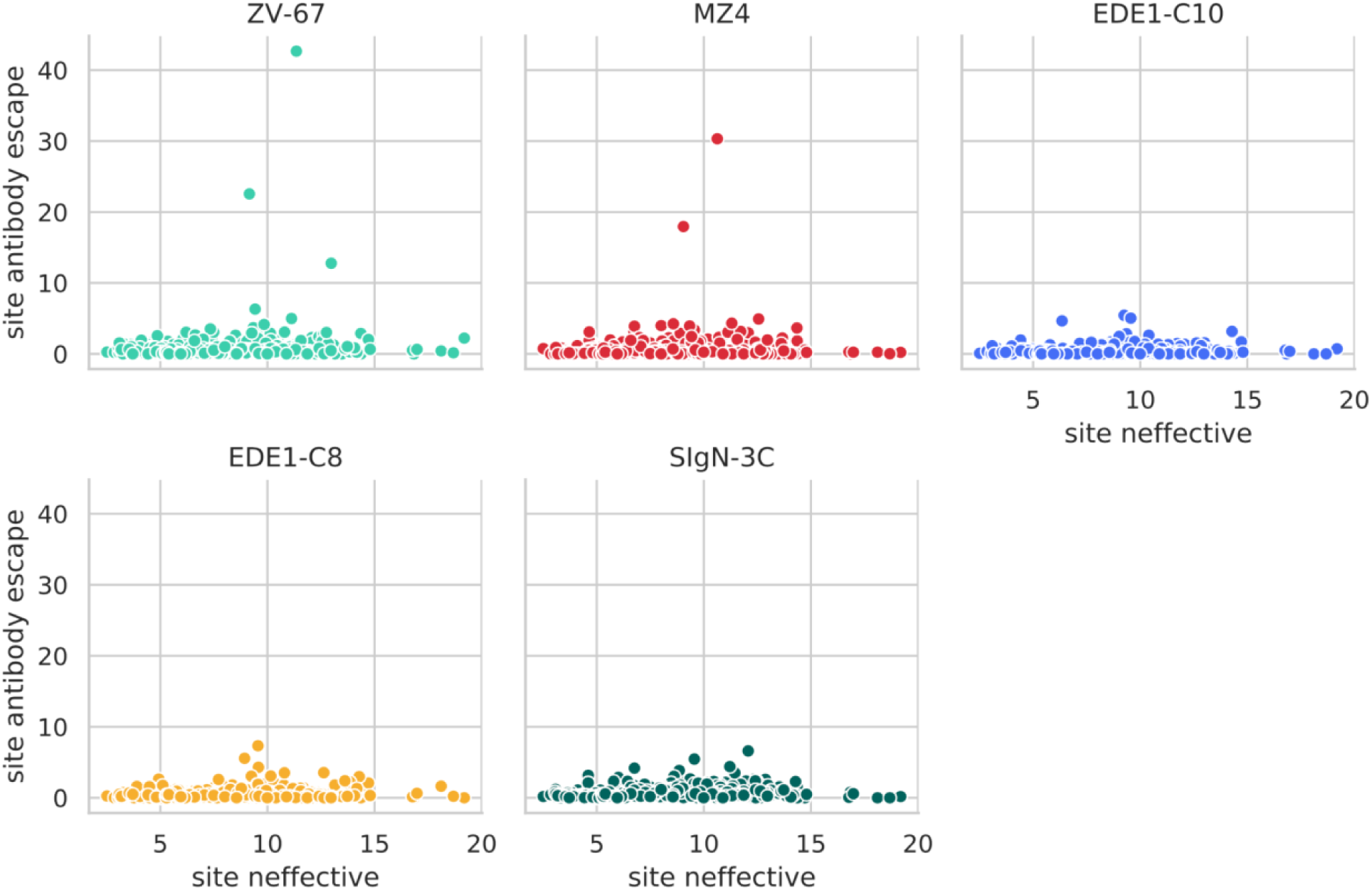
Site-level escape for all antibodies plotted against site-level neffective. Site-wise summed antibody escape is plotted against a metric of site mutational tolerance, neffective. See **Methods** for more details.

**Supplemental figure 10.**
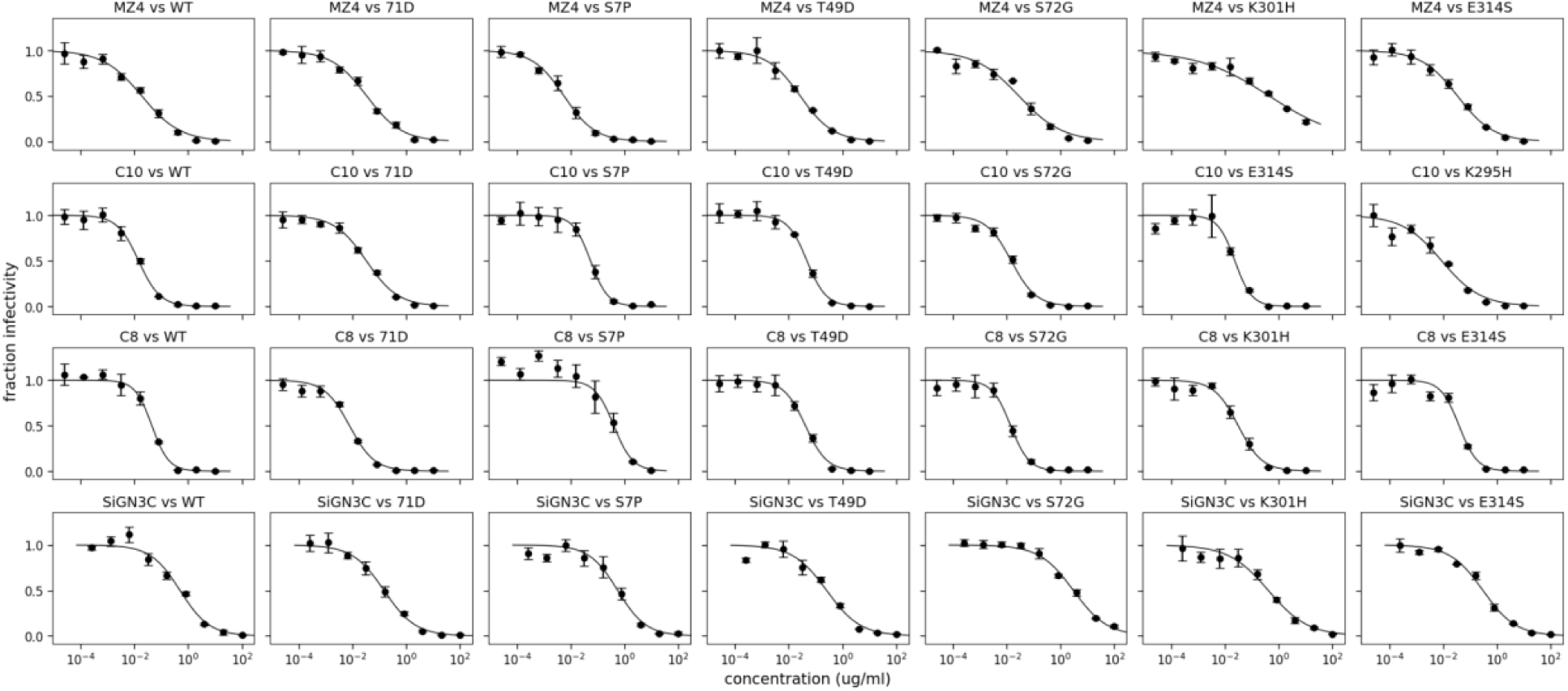
Individual Hill Curves interpolated for technical triplicate neutralization assays with Zika virus H/PF/2013. Points indicate the mean and standard error across three technical replicates.

**Supplemental figure 11.**
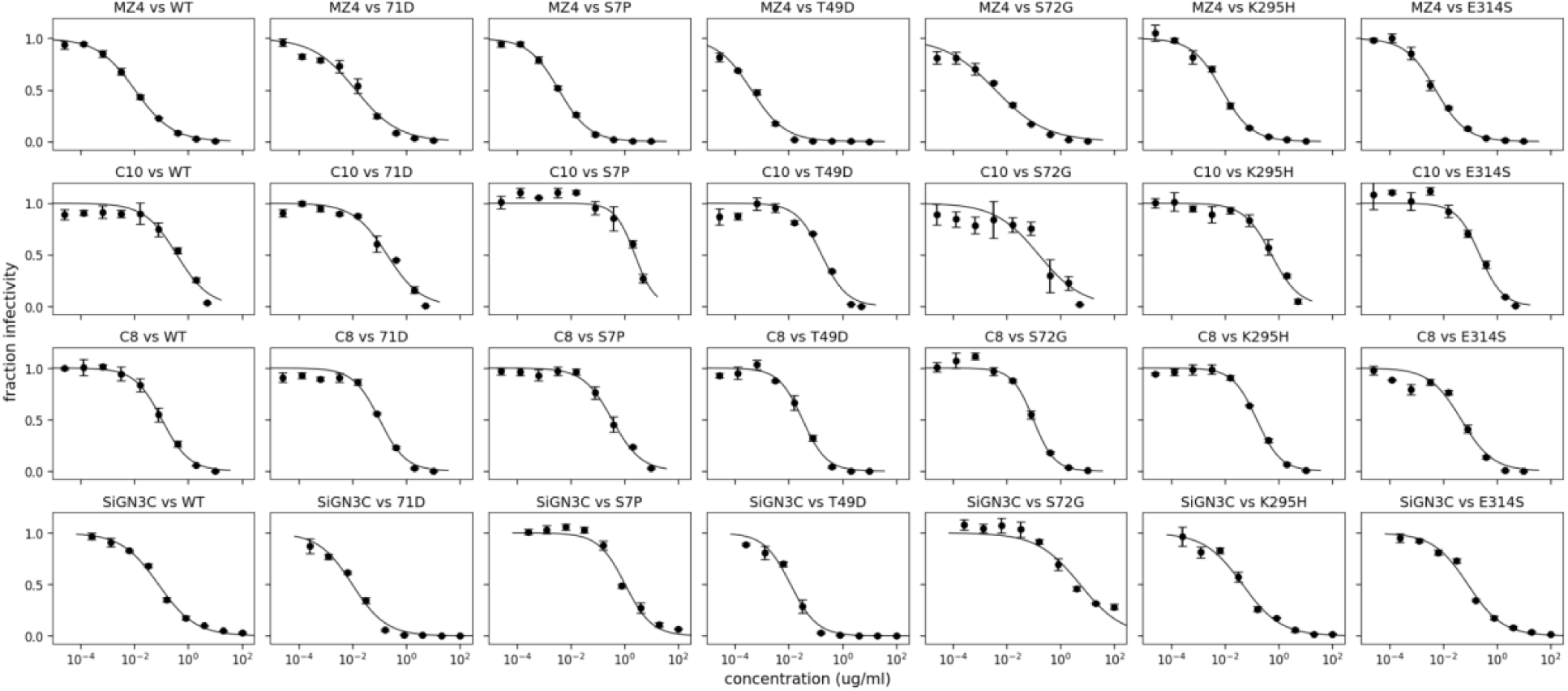
Individual Hill Curves interpolated for technical triplicate neutralization assays with dengue virus serotype 2 16681. Points indicate the mean and standard error across three technical replicates.

**Supplemental table 1.**
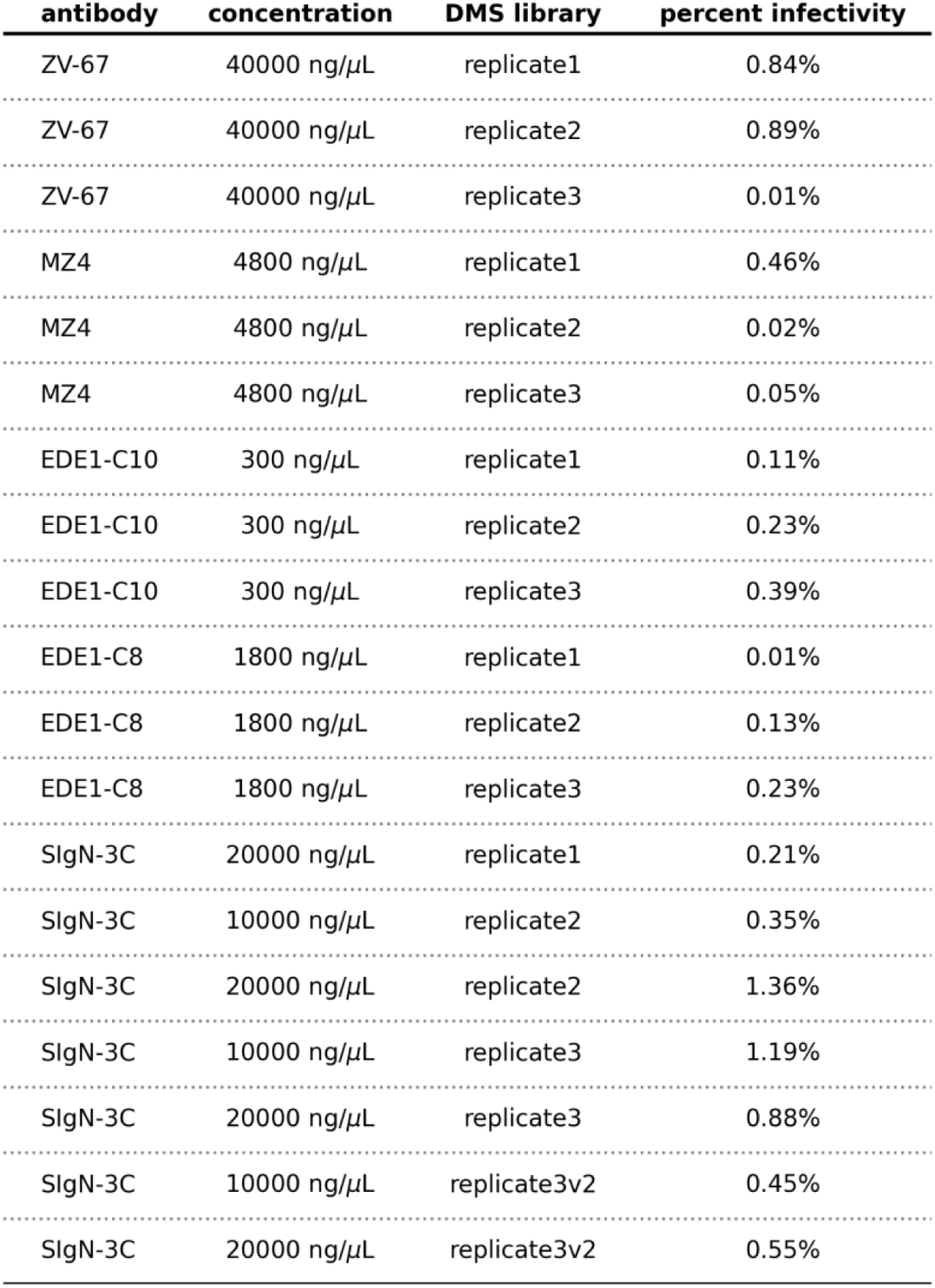
Antibody concentrations used to neutralize MR766 Zika virus DMS libraries. Antibodies were incubated at the indicated concentration with the indicated biological replicate deep mutational scanning (DMS) library. The percent infectivity was quantified by qRT-PCR. See **Methods** for details.

**Supplemental table 2.** IC50s and statistical tests for antibody ZV-67. Antibodies were incubated with viruses containing the indicated mutation created in the indicated viral background. Hill curves and IC50s were inferred in technical triplicate, and the mean and standard error (stderr) were calculated. To assess for differences between wild-type (WT) and mutant virus IC50s, a Student’s T test was performed (ttest). See **Methods** for details.

| serum | mutation | background | rep1-IC50 | rep2-IC50 | rep3-IC50 | mean-IC50 | stderr | ttest |
| --- | --- | --- | --- | --- | --- | --- | --- | --- |
| ZV-67 | WT | MR766 | 0.013 | 0.026 | 0.016 | 0.018 | 0.004 | nan |
| ZV-67 | T315Y | MR766 | 0.031 | 0.030 | 0.052 | 0.038 | 0.007 | 0.093 |
| ZV-67 | A333W | MR766 | 10.000 | 10.000 | 10.000 | 10.000 | 0.000 | **0.000** |
| ZV-67 | A333R | MR766 | 10.000 | 10.000 | 10.000 | 10.000 | 0.000 | **0.000** |
| ZV-67 | A310D | MR766 | 8.002 | 10.000 | 10.000 | 9.334 | 0.666 | **0.005** |

**Supplemental table 3.** IC50s and statistical tests for antibody MZ4. Antibodies were incubated with viruses containing the indicated mutation created in the indicated viral background. Hill curves and IC50s were inferred in technical triplicate, and the mean and standard error (stderr) were calculated. To assess for differences between wild-type (WT) and mutant virus IC50s, a Student’s T test was performed (ttest). See **Methods** for details.

| serum | mutation | background | rep1-IC50 | rep2-IC50 | rep3-IC50 | mean-IC50 | stderr | ttest |
| --- | --- | --- | --- | --- | --- | --- | --- | --- |
| MZ4 | WT | MR766 | 0.012 | 0.009 | 0.010 | 0.010 | 0.001 | nan |
| MZ4 | WT | HPF2013 | 0.018 | 0.015 | 0.022 | 0.018 | 0.002 | nan |
| MZ4 | WT | DENV2 | 0.013 | 0.007 | 0.011 | 0.011 | 0.002 | nan |
| MZ4 | V331 | MR766 | 0.008 | 0.018 | 0.014 | 0.013 | 0.003 | 0.383 |
| MZ4 | T49D | MR766 | 0.007 | 0.006 | 0.007 | 0.007 | 0.000 | **0.040** |
| MZ4 | T49D | HPF2013 | 0.014 | 0.036 | 0.026 | 0.026 | 0.006 | 0.363 |
| MZ4 | T49D | DENV2 | 0.000 | 0.000 | 0.000 | 0.000 | 0.000 | **0.026** |
| MZ4 | S7P | MR766 | 0.000 | 0.002 | 0.004 | 0.002 | 0.001 | **0.008** |
| MZ4 | S7P | HPF2013 | 0.004 | 0.011 | 0.004 | 0.006 | 0.002 | **0.018** |
| MZ4 | S7P | DENV2 | 0.004 | 0.004 | 0.003 | 0.004 | 0.000 | 0.053 |
| MZ4 | S72G | MR766 | 0.017 | 0.020 | 0.051 | 0.029 | 0.011 | 0.224 |
| MZ4 | S72G | HPF2013 | 0.019 | 0.033 | 0.030 | 0.027 | 0.004 | 0.171 |
| MZ4 | S72G | DENV2 | 0.003 | 0.005 | 0.004 | 0.004 | 0.000 | **0.047** |
| MZ4 | S368V | MR766 | 0.539 | 0.459 | 0.159 | 0.386 | 0.116 | 0.083 |
| MZ4 | S368N | MR766 | 5.861 | 4.115 | 4.950 | 4.975 | 0.504 | **0.010** |
| MZ4 | S368I | MR766 | 1.288 | 0.715 | 1.337 | 1.113 | 0.199 | **0.031** |
| MZ4 | K301H | MR766 | 2.520 | 1.498 | 1.419 | 1.812 | 0.354 | **0.037** |
| MZ4 | K301H | HPF2013 | 0.434 | 0.383 | 0.514 | 0.444 | 0.038 | **0.008** |
| MZ4 | K295H | DENV2 | 0.007 | 0.007 | 0.009 | 0.008 | 0.001 | 0.211 |
| MZ4 | G182D | MR766 | 10.000 | 10.000 | 10.000 | 10.000 | 0.000 | **0.000** |
| MZ4 | E314S | HPF2013 | 0.034 | 0.029 | 0.039 | 0.034 | 0.003 | **0.014** |
| MZ4 | E314S | DENV2 | 0.004 | 0.006 | 0.006 | 0.005 | 0.001 | 0.072 |
| MZ4 | A333W | MR766 | 0.011 | 0.014 | 0.010 | 0.012 | 0.001 | 0.447 |

**Supplemental table 4.** IC50s and statistical tests for antibody EDE1-C10. Antibodies were incubated with viruses containing the indicated mutation created in the indicated viral background. Hill curves and IC50s were inferred in technical triplicate, and the mean and standard error (stderr) were calculated. To assess for differences between wild-type (WT) and mutant virus IC50s, a Student’s T test was performed (ttest). See **Methods** for details.

| serum | mutation | background | rep1-IC50 | rep2-IC50 | rep3-IC50 | mean-IC50 | stderr | ttest |
| --- | --- | --- | --- | --- | --- | --- | --- | --- |
| EDE1-C10 | WT | MR766 | 0.012 | 0.005 | 0.008 | 0.009 | 0.002 | nan |
| EDE1-C10 | WT | HPF2013 | 0.014 | 0.014 | 0.012 | 0.013 | 0.001 | nan |
| EDE1-C10 | WT | DENV2 | 0.256 | 0.531 | 0.295 | 0.361 | 0.086 | nan |
| EDE1-C10 | V331 | MR766 | 0.003 | 0.012 | 0.008 | 0.008 | 0.003 | 0.785 |
| EDE1-C10 | T49D | MR766 | 0.023 | 0.018 | 0.019 | 0.020 | 0.002 | **0.013** |
| EDE1-C10 | T49D | HPF2013 | 0.047 | 0.057 | 0.040 | 0.048 | 0.005 | **0.017** |
| EDE1-C10 | T49D | DENV2 | 0.167 | 0.184 | 0.148 | 0.166 | 0.011 | 0.150 |
| EDE1-C10 | T315Y | MR766 | 0.009 | 0.005 | 0.013 | 0.009 | 0.002 | 0.956 |
| EDE1-C10 | T315C | MR766 | 0.006 | 0.001 | 0.012 | 0.006 | 0.003 | 0.574 |
| EDE1-C10 | S7P | MR766 | 0.010 | 0.007 | 0.021 | 0.013 | 0.004 | 0.457 |
| EDE1-C10 | S7P | HPF2013 | 0.041 | 0.082 | 0.050 | 0.057 | 0.012 | 0.070 |
| EDE1-C10 | S7P | DENV2 | 3.080 | 1.418 | 2.676 | 2.391 | 0.501 | 0.052 |
| EDE1-C10 | S72G | MR766 | 0.007 | 0.004 | 0.002 | 0.004 | 0.002 | 0.172 |
| EDE1-C10 | S72G | HPF2013 | 0.012 | 0.019 | 0.012 | 0.014 | 0.002 | 0.778 |
| EDE1-C10 | S72G | DENV2 | 0.118 | 0.019 | 0.332 | 0.156 | 0.092 | 0.181 |
| EDE1-C10 | N8T | MR766 | 0.001 | 0.006 | 0.004 | 0.004 | 0.001 | 0.132 |
| EDE1-C10 | N439K | MR766 | 0.005 | 0.005 | 0.003 | 0.004 | 0.001 | 0.139 |
| EDE1-C10 | K301H | MR766 | 0.005 | 0.004 | 0.006 | 0.005 | 0.001 | 0.202 |
| EDE1-C10 | K295H | HPF2013 | 0.010 | 0.013 | 0.004 | 0.009 | 0.003 | 0.237 |
| EDE1-C10 | K295H | DENV2 | 0.339 | 0.713 | 0.636 | 0.563 | 0.114 | 0.235 |
| EDE1-C10 | E314S | HPF2013 | 0.016 | 0.015 | 0.030 | 0.020 | 0.005 | 0.294 |
| EDE1-C10 | E314S | DENV2 | 0.168 | 0.276 | 0.252 | 0.232 | 0.033 | 0.270 |
| EDE1-C10 | A333W | MR766 | 0.004 | 0.007 | 0.005 | 0.005 | 0.001 | 0.218 |

**Supplemental table 5.** IC50s and statistical tests for antibody EDE1-C8. Antibodies were incubated with viruses containing the indicated mutation created in the indicated viral background. Hill curves and IC50s were inferred in technical triplicate, and the mean and standard error (stderr) were calculated. To assess for differences between wild-type (WT) and mutant virus IC50s, a Student’s T test was performed (ttest). See **Methods** for details.

| serum | mutation | background | rep1-IC50 | rep2-IC50 | rep3-IC50 | mean-IC50 | stderr | ttest |
| --- | --- | --- | --- | --- | --- | --- | --- | --- |
| EDE1-C8 | WT | MR766 | 0.028 | 0.020 | 0.020 | 0.022 | 0.003 | nan |
| EDE1-C8 | WT | HPF2013 | 0.038 | 0.042 | 0.057 | 0.046 | 0.006 | nan |
| EDE1-C8 | WT | DENV2 | 0.161 | 0.052 | 0.120 | 0.111 | 0.032 | nan |
| EDE1-C8 | V331 | MR766 | 0.013 | 0.015 | 0.019 | 0.016 | 0.002 | 0.123 |
| EDE1-C8 | T49D | MR766 | 0.010 | 0.025 | 0.022 | 0.019 | 0.005 | 0.537 |
| EDE1-C8 | T49D | HPF2013 | 0.061 | 0.027 | 0.039 | 0.042 | 0.010 | 0.779 |
| EDE1-C8 | T49D | DENV2 | 0.045 | 0.037 | 0.021 | 0.034 | 0.007 | 0.130 |
| EDE1-C8 | T315Y | MR766 | 0.017 | 0.042 | 0.056 | 0.038 | 0.011 | 0.292 |
| EDE1-C8 | T315C | MR766 | 0.004 | 0.026 | 0.021 | 0.017 | 0.006 | 0.493 |
| EDE1-C8 | S7P | MR766 | 0.066 | 0.060 | 0.140 | 0.089 | 0.026 | 0.120 |
| EDE1-C8 | S7P | HPF2013 | 0.598 | 0.165 | 0.351 | 0.371 | 0.126 | 0.122 |
| EDE1-C8 | S7P | DENV2 | 0.389 | 0.518 | 0.205 | 0.371 | 0.091 | 0.090 |
| EDE1-C8 | S72G | MR766 | 0.008 | 0.011 | 0.010 | 0.009 | 0.001 | **0.029** |
| EDE1-C8 | S72G | HPF2013 | 0.014 | 0.018 | 0.008 | 0.013 | 0.003 | **0.018** |
| EDE1-C8 | S72G | DENV2 | 0.108 | 0.081 | 0.105 | 0.098 | 0.009 | 0.722 |
| EDE1-C8 | N8T | MR766 | 0.026 | 0.022 | 0.013 | 0.021 | 0.004 | 0.739 |
| EDE1-C8 | N439K | MR766 | 0.004 | 0.022 | 0.017 | 0.014 | 0.005 | 0.261 |
| EDE1-C8 | K301H | MR766 | 0.010 | 0.013 | 0.007 | 0.010 | 0.002 | **0.024** |
| EDE1-C8 | K301H | HPF2013 | 0.021 | 0.053 | 0.028 | 0.034 | 0.010 | 0.366 |
| EDE1-C8 | K295H | DENV2 | 0.147 | 0.164 | 0.148 | 0.153 | 0.006 | 0.318 |
| EDE1-C8 | E314S | HPF2013 | 0.038 | 0.034 | 0.045 | 0.039 | 0.003 | 0.393 |
| EDE1-C8 | E314S | DENV2 | 0.067 | 0.031 | 0.046 | 0.048 | 0.011 | 0.177 |
| EDE1-C8 | A333W | MR766 | 0.015 | 0.007 | 0.010 | 0.011 | 0.002 | **0.032** |

**Supplemental table 6.**
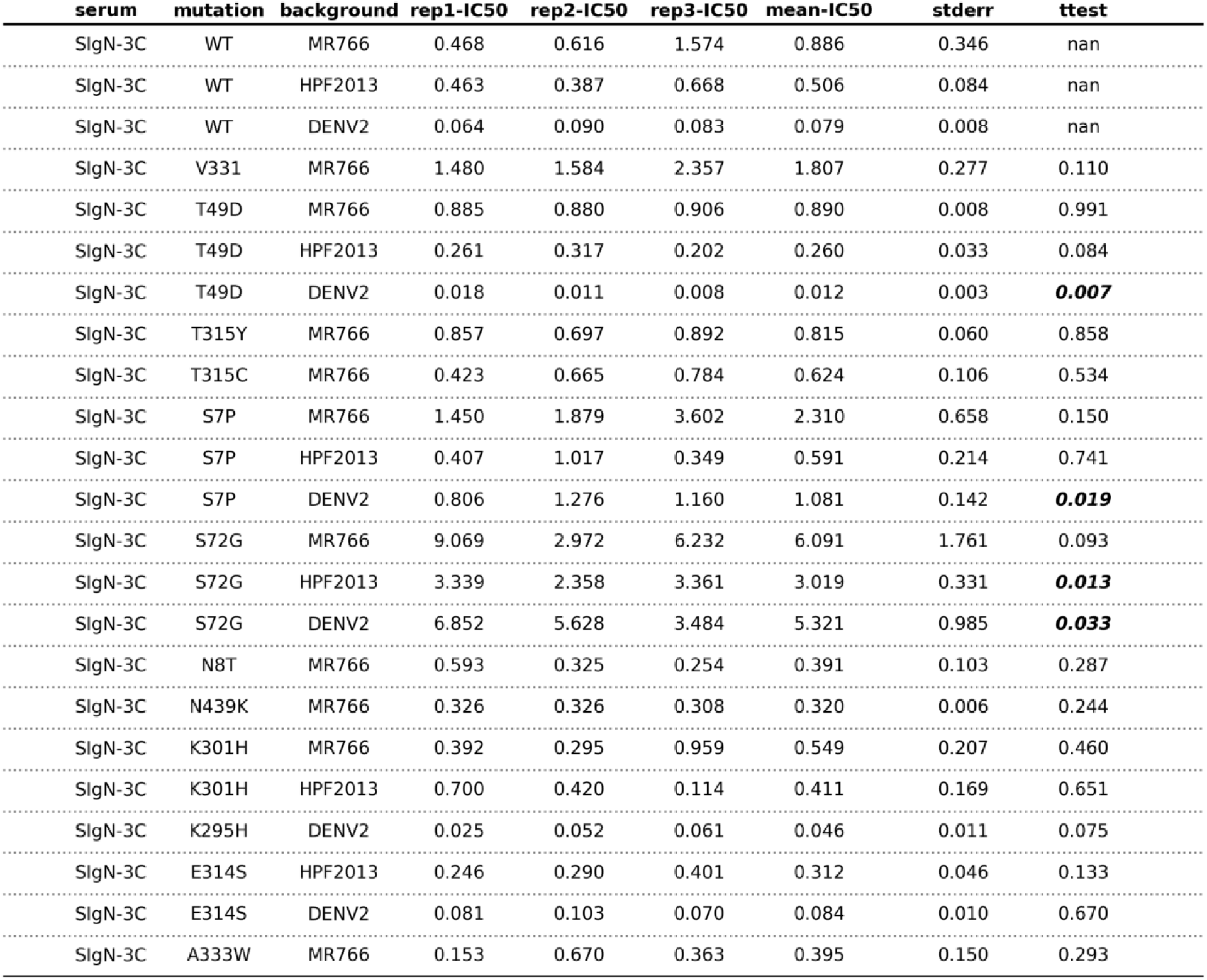
IC50s and statistical tests for antibody SIgN-3C. Antibodies were incubated with viruses containing the indicated mutation created in the indicated viral background. Hill curves and IC50s were inferred in technical triplicate, and the mean and standard error (stderr) were calculated. To assess for differences between wild-type (WT) and mutant virus IC50s, a Student’s T test was performed (ttest). See **Methods** for details.

| serum | mutation | background | rep1-IC50 | rep2-IC50 | rep3-IC50 | mean-IC50 | stderr | ttest |
| --- | --- | --- | --- | --- | --- | --- | --- | --- |
| SIgN-3C | WT | MR766 | 0.468 | 0.616 | 1.574 | 0.886 | 0.346 | nan |
| SIgN-3C | WT | HPF2013 | 0.463 | 0.387 | 0.668 | 0.506 | 0.084 | nan |
| SIgN-3C | WT | DENV2 | 0.064 | 0.090 | 0.083 | 0.079 | 0.008 | nan |
| SIgN-3C | V331 | MR766 | 1.480 | 1.584 | 2.357 | 1.807 | 0.277 | 0.110 |
| SIgN-3C | T49D | MR766 | 0.885 | 0.880 | 0.906 | 0.890 | 0.008 | 0.991 |
| SIgN-3C | T49D | HPF2013 | 0.261 | 0.317 | 0.202 | 0.260 | 0.033 | 0.084 |
| SIgN-3C | T49D | DENV2 | 0.018 | 0.011 | 0.008 | 0.012 | 0.003 | **0.007** |
| SIgN-3C | T315Y | MR766 | 0.857 | 0.697 | 0.892 | 0.815 | 0.060 | 0.858 |
| SIgN-3C | T315C | MR766 | 0.423 | 0.665 | 0.784 | 0.624 | 0.106 | 0.534 |
| SIgN-3C | S7P | MR766 | 1.450 | 1.879 | 3.602 | 2.310 | 0.658 | 0.150 |
| SIgN-3C | S7P | HPF2013 | 0.407 | 1.017 | 0.349 | 0.591 | 0.214 | 0.741 |
| SIgN-3C | S7P | DENV2 | 0.806 | 1.276 | 1.160 | 1.081 | 0.142 | **0.019** |
| SIgN-3C | S72G | MR766 | 9.069 | 2.972 | 6.232 | 6.091 | 1.761 | 0.093 |
| SIgN-3C | S72G | HPF2013 | 3.339 | 2.358 | 3.361 | 3.019 | 0.331 | **0.013** |
| SIgN-3C | S72G | DENV2 | 6.852 | 5.628 | 3.484 | 5.321 | 0.985 | **0.033** |
| SIgN-3C | N8T | MR766 | 0.593 | 0.325 | 0.254 | 0.391 | 0.103 | 0.287 |
| SIgN-3C | N439K | MR766 | 0.326 | 0.326 | 0.308 | 0.320 | 0.006 | 0.244 |
| SIgN-3C | K301H | MR766 | 0.392 | 0.295 | 0.959 | 0.549 | 0.207 | 0.460 |
| SIgN-3C | K301H | HPF2013 | 0.700 | 0.420 | 0.114 | 0.411 | 0.169 | 0.651 |
| SIgN-3C | K295H | DENV2 | 0.025 | 0.052 | 0.061 | 0.046 | 0.011 | 0.075 |
| SIgN-3C | E314S | HPF2013 | 0.246 | 0.290 | 0.401 | 0.312 | 0.046 | 0.133 |
| SIgN-3C | E314S | DENV2 | 0.081 | 0.103 | 0.070 | 0.084 | 0.010 | 0.670 |
| SIgN-3C | A333W | MR766 | 0.153 | 0.670 | 0.363 | 0.395 | 0.150 | 0.293 |

## Methods

### Cell lines

293T cells (Cat # CRL-11268, ATCC, Manassas, VA) and Vero-C1008 cells (Cat # CRL-1586, ATCC, Manassas, VA) were cultured in Dulbecco’s modified Eagle’s medium (DMEM) (Gibco BRI; Life Technologies, Gaithersburg, MD) supplemented with 7% fetal bovine serum (FBS) (Gibco BRI; Life Technologies, Gaithersburg, MD) and 100 U/mL penicillin-streptomycin (Gibco BRI; Life Technologies, Gaithersburg, MD). Raji cells stably expressing dendritic cell-specific intercellular adhesion molecule-grabbing nonintegrin-related (Raji/DC-SIGNR)^45^ (provided by Theodore C. Pierson, National Institutes of Health) were maintained in Roswell Park Memorial Institute (RPMI) 1640 supplemented with GlutaMAX (Gibco BRI; Life Technologies, Gaithersburg, MD), 7% FBS and 100 U/mL penicillin-streptomycin. All cell lines were maintained at 37 °C in 5% CO2.

### Antibodies

EDE1-C10, EDE1-C8 and SIgN-3C are human broadly neutralizing monoclonal antibodies targeting a complex quaternary E protein epitope, and were produced as IgG1 as previously described^17^. MZ4 heavy and light chain expression plasmids^20^ (provided by Shelly Krebs, Walter Reed Army Institute of Research) were used to produce IgG1 antibodies targeting a domain III epitope on E protein as previously described^17^. ZV-67 is a mouse monoclonal targeting Zika virus E protein^33^ (provided by Michael S. Diamond, Washington University in St. Louis)

### E protein alignment and phylogenetic tree inference

We selected a single representative strain of each dengue serotype 1-4, two strains of Zika virus, and a strain of West Nile virus as an outgroup. We downloaded both protein and DNA sequences for each of these strains from NCBI and used `mafft`^46^ to align protein sequences. This protein alignment was used to compare Zika virus and dengue virus sequences for single mutant pseudovirus particle generation in **Figure 5**. We next inferred a phylogenetic tree using IQ-Tree^47^. We selected a protein sequence model (LG+I+G4^48,49^) with the lowest BIC from IQ-Tree’s ModelFinder^50^, but compared similar models and found they produced qualitatively similar trees. Trees were then rendered using ETE3^51^ (https://github.com/jbloomlab/ZIKV_MAP_GooLab/blob/main/paper_figures/flavivirus_alignment/ <u>draw_phylogeny_from_newick.ipynb</u>).

### Zika virus mutant E protein virus libraries

These libraries were designed and described in Sourisseau et al^34^. Briefly, these libraries were generated by tiling codon mutagenesis of Zika virus MR766 E gene in full biological triplicate, resulting in slightly different library compositions at the nucleotide and amino acid level^34^. Using a reverse genetics system^52^, these mutant plasmid libraries are transfected into cells, and 3 days later, harvested to produce mutant virus libraries. To ensure the viral variant phenotype is matched to its own packaged genome, thus purging the libraries of unfit E variants, the virus libraries were passaged at low multiplicity of infection on Vero cells. These genotype-phenotype matched libraries were then titrated on Vero cells to determine infectious units per mL.

### Deep mutational scanning

Deep mutational scanning was performed as described^34,35,53^. Briefly, virus libraries were diluted to 1 million infectious units per mL in DMEM supplemented with 2% FBS and incubated with antibodies diluted to the concentrations indicated in **Supplemental Table 1** for 1 hour at 37°C. The entire antibody-virus library mixtures were then added to Vero cells and cultured for 24 hours at 37°C. Cells were then washed extensively with PBS before lysis for RNA extraction. Viral cDNA was reverse-transcribed and PCR amplified as described previously^34^. Each antibody was profiled with each biological triplicate mutant virus library. In addition, one antibody, SIgN-3C was profiled in technical duplicate. To increase fidelity of Illumina deep sequencing, we used the same barcoded-subamplicon sequencing strategy as reported in Doud and Bloom^54^. The primers and details of this protocol all match those described in Sourisseau et al^34^.

### Determining percent infectivity with Zika virus qRT-PCR

To determine the percent infectivity reported in **Supplemental Table 1** we used qRT-PCR against Zika virus E gene (primers reported in Sourisseau et al.^34^) and GAPDH (primers reported in Zhang et al.^55^). To create a standard curve of infectivity by the mutant virus libraries, we made 10-fold dilutions of the mutant virus library and infected cells with these standards in technical duplicate. This standard curve was generated independently for each deep mutational scanning experiment. Total RNA was extracted from both virus-antibody-infected cells and virus standard curve-infected cells for quantitative qRT-PCR using the One-Step RT-qPCR Kit (Quanta Bio). We then fit a linear regression to the difference between the E and GAPDH standard curve Ct values and used this regression to estimate the percent infectivity of the experimental samples.

### Analysis and visualization of deep sequencing data

The deep sequencing data were analyzed and visualized using `dms_tools2` (https://jbloomlab.github.io/dms_tools2/)^56,57^. All code and documentation are available online (https://github.com/jbloomlab/ZIKV_MAP_GooLab). Methods used in this paper include calculation of differential selection (`dms2_batch_diffsel`), which was used as our metric of antibody escape in **Figures 2B, 3 and 5** and **Supplemental Figures 2-6**. Differential selection was calculated independently for biological replicate antibody selections, and the median values are shown. In **Figure 3**, zoomed logo plots were produced using custom code and the Python package `dmslogo` (https://jbloomlab.github.io/dmslogo/). For estimation of mutation-level tolerance in Figure 5A, we calculated the log ratio of mutant to wild-type amino acid preferences (`dms2_batch_prefs`). For estimation of site-level mutational flexibility in **Figure 5B** and **Supplemental Figure 9**, we used the metrics entropy and neffective, respectively (`dms_tools2.prefs.prefsEntropy`).

Protein structures of Zika virus E protein and all antibodies were downloaded from Protein Data Bank. To project summed site-level escape on E protein surface, b-factors were redefined using the function `reassign_b_factors` from `polyclonal` (https://jbloomlab.github.io/polyclonal/)^58^. These modified files were then modeled using The PyMOL Molecular Graphics System, Version 2.0 Schrödinger, LLC.

### Validation of deep mutational scanning using reporter virus particles

Reporter virus particles were generated as previously described^17^. Briefly, a plasmid containing West Nile virus non-structural proteins and a green fluorescent protein (GFP) reporter (so-called ‘replicon’ plasmid)^59^ was co-transfected into 293T cells with a plasmid containing flaviviral structural capsid, pre-M and E proteins (so-called ‘CprME’ plasmid). The CprME plasmids used were: Zika virus (MR766)^1^, Zika virus (H/PF/2013)^1^ and dengue serotype 2 (16681)^60^. Together, 1 µg of replicon was added to 3 µg of CprME plasmid and subsequently mixed with Lipofectamine 3000 (Invitrogen, Waltham, MA) and OptiMEM (Gibco BRI; Life Technologies, Gaithersburg, MD) before being added to 293T cells. After incubation at 37°C for 4-6 hours, the transfection mixture was replaced with low-glucose DMEM supplemented with 7% FBS (Gibco BRI; Life Technologies, Gaithersburg, MD). Cells were then cultured for 4-6 days at 30°C and 5% CO^2^. Subsequently, reporter virus particle-containing supernatant was passed through a 0.22 µm Millex SteriFlip Vacuum-driven Filtration System (Sigma-Aldrich, Burlington, MA) before aliquoting and storage at −80°C. Infectious units per ml were determined by titration of reporter virus particles onto Raji/DC-SIGNR cells and then incubated at 37°C for 48 hours. Cells were fixed in 2% paraformaldehyde (Electron Microscopy Sciences, Hatfield, PA) before percent of GFP-positive cells was determined by high-throughput flow cytometry (Intellicyt iQue Screener PLUS, Sartorius, Gottingen, Germany).

Single mutant reporter virus particles were generated by site-directed mutagenesis of the CprME plasmid. As in previous work^61^, PCR reactions were run using partially overlapping primer pairs containing the desired mutation, followed by HiFi assembly and sequence verification by Plasmidsaurus. These mutated CprME plasmids were then used to generate single-mutant reporter virus particles as above.

For dose-dependent neutralization assays with reporter virus particles, antibodies of interest were diluted to an initial concentration of 10-50 µg/mL in RPMI supplemented with 7% FBS and then 5-fold serially diluted across a 384-well v-shape microplate (Grenier Bio-One, Kremsmünster, Austria). Reporter virus particles were diluted to 5-10% expected infectivity and added to diluted antibodies. Antibody-virus mixtures were incubated at 37°C for 4-6 hours. Next, 20 µL of 1e6/mL Raji/DC-SIGNR cells were added and the mixture was incubated at 37°C for 48 hours. As above, cells were fixed in 2% paraformaldehyde prior to measurement of %GFP by high-throughput flow cytometry. Neutralization assays were performed in technical triplicate on the same day to minimize batch effects.

Downstream analysis was performed using custom Python code with mutant viruses (https://github.com/jbloomlab/ZIKV_MAP_GooLab/tree/main/paper_figures/neutralizations_esca pe_mutants) and wild-type viruses (https://github.com/jbloomlab/ZIKV_MAP_GooLab/tree/main/paper_figures/neutralizations_WT). Fraction infectivity was calculated by dividing samples containing antibodies by the average of those containing no antibody. Plotting of neutralization curves and interpolation of Hill curves and IC50s were performed using the Python package `neutcurve` (https://jbloomlab.github.io/neutcurve/). Tables of percent infectivty, replicate IC50s and statistical tests in **Supplemental Tables 1-6** were rendered with custom code (https://github.com/jbloomlab/ZIKV_MAP_GooLab/blob/main/paper_figures/neutralizations_esca pe_mutants/all_nabs_neutcurve.ipynb) using the Python package `pyMSAviz` (https://moshi4.github.io/pyMSAviz/). Statistical tests used and more details can be found in the above linked directories and corresponding Jupyter notebooks.

## Data availability

The computer code used to analyze deep sequencing data can be found on GitHub (https://github.com/jbloomlab/ZIKV_MAP_GooLab). All raw sequencing data was deposited on the Sequence Read Archive under BioProject accession number PRJNA530795 and BioSample accession number SAMN37316695.

